# BRCA2 promotes genomic integrity and therapy resistance primarily through its role in homology-directed repair

**DOI:** 10.1101/2023.04.11.536470

**Authors:** Pei Xin Lim, Mahdia Zaman, Maria Jasin

## Abstract

**Highlights:**

- Gap suppression requires BRCA2 C-terminal RAD51 binding in mouse and human cells
- *Brca2* heterozygosity in mice results in fork protection and gap suppression defects
- Gap suppression mitigates sensitivity to hmdU, but only when HDR is unperturbed
- HDR deficiency is the primary driver of chemotherapeutic sensitivity

**eTOC blurb:** Lim *et al*. report that gap suppression as well as fork protection require BRCA2 stabilization of RAD51 filaments in human and mouse cells but have minimal impact on genome integrity, oncogenesis, and drug resistance. BRCA2 suppression of PRIMPOL-mediated replication gaps confers resistance to the nucleotide hmdU, incorporation of which leads to cytotoxic abasic sites.This effect is diminished when HDR is abrogated.

**Summary:** Tumor suppressor BRCA2 functions in homology-directed repair (HDR), protection of stalled replication forks, and suppression of replicative gaps. The relative contributions of these pathways to genome integrity and chemotherapy response are under scrutiny. Here, we report that mouse and human cells require a RAD51 filament stabilization motif in BRCA2 for both fork protection and gap suppression, but not HDR. Loss of fork protection and gap suppression do not compromise genome instability or shorten tumor latency in mice or cause replication stress in human mammary cells. By contrast, HDR deficiency increases spontaneous and replication stress-induced chromosome aberrations and tumor predisposition. Unlike with HDR, fork protection and gap suppression defects are also observed in *Brca2* heterozygous mouse cells, likely due to reduced RAD51 stabilization at stalled forks and gaps. Gaps arise from PRIMPOL activity, which is associated with sensitivity to 5-hydroxymethyl-2’-deoxyuridine due to the formation of abasic sites by SMUG1 glycosylase and is exacerbated by poly(ADP-ribose) polymerase inhibition. However, HDR deficiency ultimately modulates sensitivity to chemotherapeutics, including PARP inhibitors.

## Introduction

Genomic instability is one of the hallmarks of tumorigenesis (Hanahan and Weinberg, 2011). The tumor suppressor BRCA2 is essential for the maintenance of genomic integrity, presumably through the suppression of endogenous replication stress (Feng and Jasin, 2017), a characteristic of pre-cancerous lesions (Macheret and Halazonetis, 2015). Concordantly, carriers of *BRCA2* germline mutations are predisposed to various cancers, such as breast, ovarian, and prostate, with tumors typically exhibiting loss of heterozygosity (Chen *et al*., 2018; Maxwell *et al*., 2017; Pennington *et al*., 2014; Pritchard *et al*., 2016; Skoulidis *et al*., 2010; Tavtigian *et al*., 1996; Wooster *et al*., 1995). These tumors have large numbers of chromosome aberrations as well as single base and indel mutations and can be targeted by specific cancer chemotherapeutics (Davies *et al*., 2017).

Mechanistically, BRCA2 is critical for the error-free repair of DNA double-strand breaks (DSBs) by homologous recombination, also termed homology-directed repair (HDR) (Moynahan *et al*., 2001). During HDR, DNA end resection proteins process DSBs to create single-stranded DNA (ssDNA) onto which BRCA2 nucleates RAD51 filaments (Prakash *et al*., 2015; Zhao *et al*., 2019). The RAD51 nucleoprotein filament invades homologous DNA to prime repair synthesis and restore the original sequence prior to DNA damage or, in the case of replication stress, to restore the integrity of the replication fork (Scully *et al*., 2019).

BRCA2 interacts with RAD51 through eight BRC motifs in the central part of the protein to promote RAD51 filament formation, and a distinct, conserved site at the C terminus to stabilize RAD51 filaments (Prakash *et al*., 2015). Murine *Brca2* truncations that completely abrogate the BRC repeats lead to embryonic lethality, whereas truncations that maintain some of the BRC repeats allow a few mice to survive to term, although these mice have a short lifespan due to highly penetrant tumor formation (Connor *et al*., 1997; Friedman *et al*., 1998; Moynahan, 2002). By contrast, C-terminal truncation is compatible with survival but predisposes to late onset tumors (Donoho *et al*., 2003; McAllister *et al*., 2002). Cells from these truncations are hypersensitive to genotoxins and demonstrate genomic instability and, where tested, HDR defects (Kass *et al*., 2016; Moynahan *et al*., 2001; Navarro *et al*., 2006). However, BRCA2 with a point mutation at the C-terminal RAD51 interaction site substantially restores HDR to BRCA2-deficient hamster and human cells (Feng and Jasin, 2017; Schlacher *et al*., 2011; Siaud *et al*., 2011), indicating that it is not critical for HDR.

The role of BRCA2 is not limited to HDR. BRCA2 is also important for protecting stalled replication forks (Schlacher *et al*., 2011) and for suppressing the formation of ssDNA gaps during replication (Panzarino *et al*., 2021; Quinet *et al*., 2020). When a replication fork stalls due to the depletion of free nucleotides or DNA lesion blockade, chromatin remodeling proteins can promote fork reversal to slow fork progression (Bai *et al*., 2020; Dungrawala *et al*., 2017; Taglialatela *et al*., 2017; Vujanovic *et al*., 2017; Zellweger *et al*., 2015). In BRCA2 mutants, the nascent strands at the reversed fork are degraded by MRE11 and other nucleases (Hashimoto *et al*., 2010; Kolinjivadi *et al*., 2017; Mijic *et al*., 2017; Schlacher *et al*., 2011). In contrast to HDR, mutation of the C-terminal RAD51 interaction site also leads to fork protection defects (Feng and Jasin, 2017; Schlacher *et al*., 2011), implying that the reversed fork is normally protected by BRCA2 C-terminal stabilization of RAD51 filaments.

Alternatively, repriming of DNA replication can occur downstream of stalled forks by the primase-polymerase PRIMPOL (Bai *et al*., 2020; Mouron *et al*., 2013; Quinet *et al*., 2020). However, repriming DNA creates ssDNA gaps that must be protected and filled in to maintain the genome. The role of BRCA2 in gap suppression is two-fold: direct inhibition of PRIMPOL activity through its interaction with MCM10 (Kang *et al*., 2021) and inhibition of MRE11-mediated degradation of the ssDNA gaps during fill in by translesion polymerases (Tirman *et al*., 2021), although the domain that dictates the gap suppression function of BRCA2 is unknown. Thus, BRCA2 is involved in multiple genome maintenance pathways: HDR, fork protection, and gap suppression.

To investigate which function(s) of BRCA2 are important for genomic stability, tumor suppression, and therapy response, we generated a mouse model with a point mutation that disrupts the C-terminal RAD51 interaction site, S3214A (abbreviated hereafter as SA). As with the cognate human mutation, this SA mutation does not impair HDR but renders stalled replication forks vulnerable to MRE11-mediated nucleolytic degradation. Surprisingly, disruption of the C-terminal RAD51 interaction site also increases PRIMPOL-mediated ssDNA gap accumulation, which was also confirmed in human cells. Unlike HDR, fork protection and gap suppression are also impaired in heterozygous mutant cells, implying that these processes are sensitive to gene dosage, consistent with a stabilization, rather than catalytic, role for BRCA2. Cells specifically deficient in gap suppression are sensitive to exogenous 5-hydroxymethyl-2’-deoxyuridine (hmdU), which can be rescued by PRIMPOL knockdown or removal of SMUG1, which creates abasic sites at hmdU sites in the genome.

These results affirm PRIMPOL function in repriming DNA synthesis downstream of stalled forks encountering abasic sites. Unlike HDR-deficient mice, the fork protection/gap suppression-deficient mice were not predisposed to tumorigenesis and they had minimal spontaneous and induced chromosome aberrations.

Further, sensitivity to chemotherapeutics, including the poly(ADP-ribose) inhibitor olaparib, was highest in HDR-deficient cells. Thus, while BRCA2 plays crucial roles in three separable pathways, HDR is the most important function in maintaining genome integrity.

## Results

### *Brca2^SA/SA^* mice are viable

Alanine or glutamate substitution at serine 3291 of the human BRCA2 gene, which abrogates RAD51 interaction in peptide studies (Esashi *et al*., 2005), leads to a marked deficiency in the protection of stalled replication forks from MRE11-dependent nucleolytic degradation (Feng and Jasin, 2017; Schlacher *et al*., 2011). To understand the physiological consequences of impaired fork protection, we introduced an alanine substitution at the cognate amino acid 3214 in the mouse *Brca2* gene, generating the *Brca2^S3214A^* (*Brca2^SA^*) allele (**Figure 1A**). The mutation was introduced in fertilized mouse eggs using TALEN-mediated gene targeting and a donor DNA fragment that also included four silent mutations to make the donor resistant to TALEN-mediated cleavage and to facilitate subsequent genotyping (**Figures S1A-S1C**). We evaluated the *Brca2^SA^* allele in tandem with a second *Brca2* allele, *Brca2^Δ27^*, which has a 190 amino acid terminal deletion starting at position 3139 (McAllister *et al*., 2002) (**Figure 1A**). Homozygosity of this *Δ27* allele has been shown to cause defects in both replication fork protection (Schlacher *et al*., 2011) and HDR (Kass *et al*., 2016). Upon intercrossing *Brca2^+/SA^* animals, *Brca2^SA/SA^* offspring were obtained at the expected Mendelian ratio (∼25%) (**Figure S1D**). By contrast, *Brca2^Δ27/Δ27^* mice were obtained at a sub-Mendelian ratio (**Figure S1D**, ∼12% versus 25%), as previously reported (McAllister *et al*., 2002), indicating developmental issues during embryogenesis.

**Figure 1.**
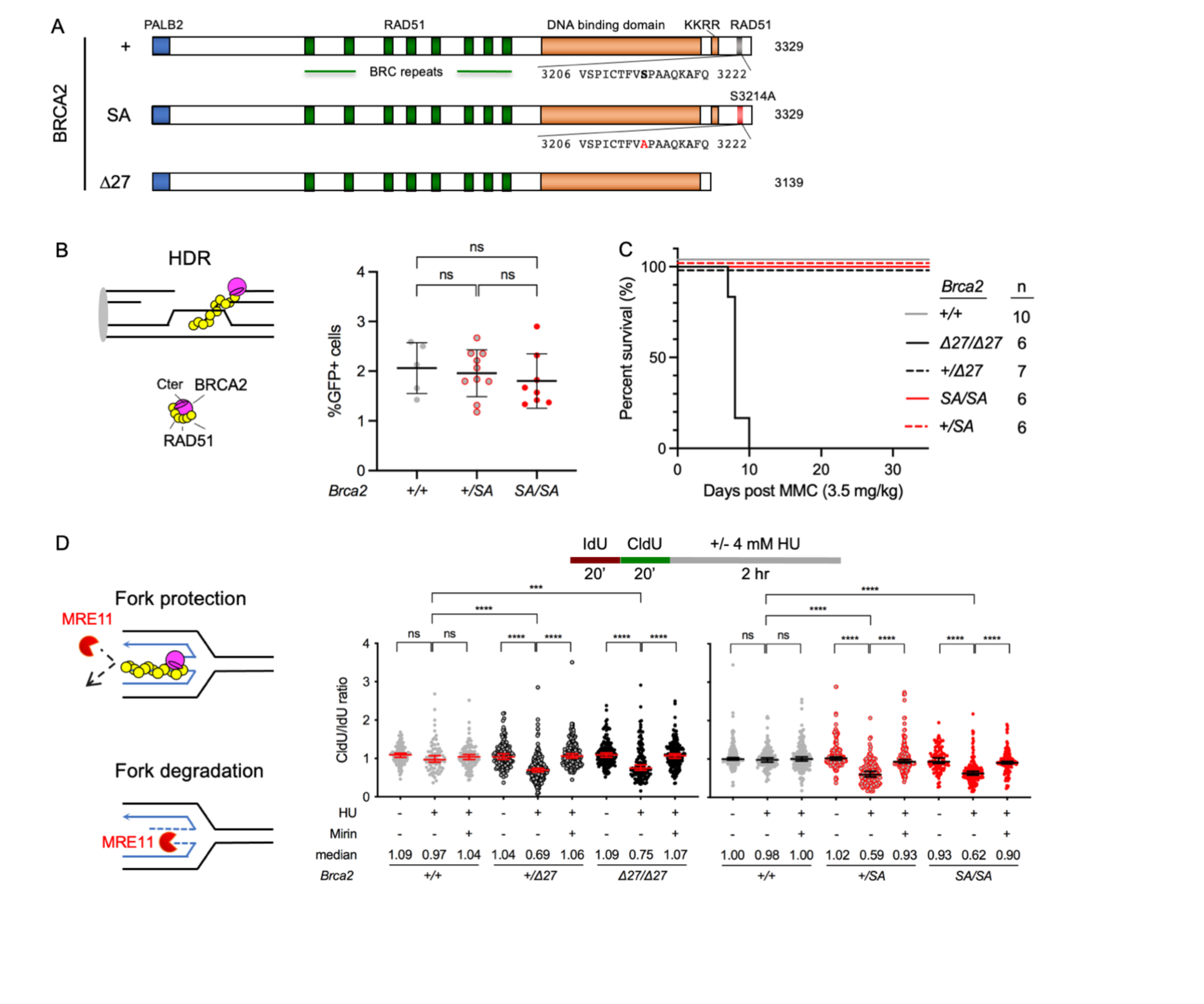
BRCA2 C-terminal RAD51-interacting site is dispensable for HDR but crucial for stalled fork protection in mouse cells. **A.** Murine BRCA2 mutant domain structures. Murine BRCA2 protein is 3329 amino acids and interacts with RAD51 at the BRC repeats and at a distinct site in the C terminus that is mutated in BRCA2 S3214A (abbreviated “SA”). BRCA2 Δ27 is a truncation of 190 amino acids that deletes the C-terminal RAD51 interacting site, as well as a recently identified DNA binding motif KKRR (Kwon *et al*., 2023). **B.** HDR is not significantly affected in *Brca2^SA/SA^* mouse cells. The diagram depicts BRCA2 nucleation of RAD51 nucleoprotein filaments to promote the strand invasion step of HDR. HDR was analyzed using an integrated DR-GFP reporter in primary ear fibroblasts. Statistical analysis, unpaired Student’s t test (mean ± 1 SD). ns, not significant. **C.** *Brca2^Δ27/Δ27^* mice, but not *Brca2^SA/SA^* mice, are sensitive to the crosslinking agent mitomycin C (MMC). The Kaplan-Meier survival curve of *Brca2* mice after intraperitoneal injection of 3.5 mg/kg MMC is shown. **D.** Stalled fork degradation in *Brca2* homozygous and heterozygous cells. Diagram shows BRCA2 and RAD51 protecting a reversed fork from MRE11-mediated nucleolytic attack. Primary MEFs were pulsed with IdU and CldU sequentially for 20 min and then incubated with or without 4 mM HU for 2 hr. Cells were treated with 50 µM mirin where indicated throughout the timeline. For each condition, at least 200 individual DNA fibers were counted. Statistical analysis, Mann-Whitney U test (median ± 95% confidence intervals). ns, not significant; ***, p<0.001; ****, p<0.0001. See Figure S2A,B for replicates.

### *Brca2^SA/SA^* cells and mice are HDR proficient

Previous studies have shown that introduction of BRCA2 S3291A into BRCA2-deficient hamster cells restores HDR proficiency to wild-type BRCA2 levels (Schlacher *et al*., 2011), and similarly that S3291A or S2391E expression in human cell lines leads to only mild to moderate HDR defects (Feng and Jasin, 2017; Rickman *et al*., 2020). To determine the impact of the *SA* allele in mice, we derived isogenic lines of primary ear fibroblasts from adult mice carrying the *Pim1^DR-GFP^* allele (Kass *et al*., 2013) and then infected these cells with lentivirus for I-SceI-expression to induce HDR (Prakash *et al*., 2021). No significant difference in the percentage of GFP-positive cells was observed among *Brca2^+/+^*, *Brca2^+/SA^* and *Brca2^SA/SA^* ear fibroblasts (averages of 2.06%, 1.96% and 1.80%, respectively, **Figure 1B**). This contrasts with *Brca2^Δ27/Δ27^* ear fibroblasts, which have a reported 11-fold reduction in HDR (Kass *et al*., 2016). Total I-SceI site loss was between 11% to 13%, indicating that the overall efficiencies of double-strand break induction and repair were similar across genotypes (**Figure S1E**).

HDR mutants are typically sensitive to DNA crosslinking agents. To further confirm that the *SA* allele does not affect HDR, we performed mitomycin C (MMC) sensitivity experiments in both *Δ27* and *SA* mutant mice. MMC was injected intraperitoneally (3.5 mg/kg), and mice were monitored for survival. *Brca2^Δ27/Δ27^* mice died within 10 days after injection, whereas all *Brca2^SA/SA^* mice survived well past 30 days, similar to control mice (**Figure 1C**). Thus, the RAD51 interaction site within the BRCA2 C-terminal domain appears to be dispensable for HDR in murine cells and in mice.

### Stalled fork protection defects with *Brca2^SA^* and *Brca2^Δ27^* homo- or heterozygosity

An intact C-terminal RAD51 interaction site has previously been shown to be crucial for the protection of nascent DNA at stalled forks in human and rodent cell lines (Feng and Jasin, 2017; Rickman *et al*., 2020; Schlacher *et al*., 2011). To confirm this in primary mouse cells, we derived isogenic lines of primary murine embryonic fibroblasts (MEFs), pulse-labeled the cells successively with 5-iodo-2’-deoxyuridine (IdU) and 5- chloro-2’-deoxyuridine (CldU), followed by treatment with hydroxyurea (HU) to impair fork progression (**Figures 1D, S2A and S2B**). As expected, HU-treated *Brca2^Δ27/Δ27^*and *Brca2^SA/SA^* primary MEFs had significantly reduced CldU/IdU tract length ratios in comparison to untreated cells and HU-treated *Brca2^+/+^*primary MEFs cells, indicating the degradation of nascent strands at stalled forks in these mutants. Importantly, this defect can be rescued by treatment with the MRE11 inhibitor mirin, consistent with previous observations that the C-terminal RAD51-interaction site is required to protect stalled forks from MRE11-dependent nucleolytic degradation (Schlacher *et al*., 2011).

Recent reports of defects in replication fork protection in heterozygous mutants of *Brca1* and *Bard1* (Billing *et al*., 2018; Pathania *et al*., 2014) prompted us to also examine *Brca2* heterozygous cells. Both *Brca2^+/Δ27^*and *Brca2^+/SA^* primary MEFs displayed reduced CldU/IdU ratios, and this was rescued by mirin treatment (**Figures 1D, S2A and S2B**). Thus, replication fork protection requires homozygosity for an intact BRCA2 C-terminal RAD51 interaction.

### Impaired gap suppression with *Brca2^SA^* and *Brca2^Δ27^* homo- or heterozygosity

In addition to protecting forks, cells under replication stress can reprime DNA synthesis ahead of stalled forks to continue replication, creating ssDNA gaps that are sensitive to S1 nuclease digestion (Berti *et al*., 2020; Quinet *et al*., 2020). To determine whether the BRCA2 C-terminal RAD51-interaction site is important for gap suppression, we pulse-labeled primary MEFs with IdU, followed by simultaneous CldU labeling and low dose HU treatment (**Figures 2A, S2C and S2D**). Remarkably, both *Brca2^Δ27/Δ27^* and *Brca2^SA/SA^* MEFs showed reduced CldU/IdU ratios upon S1 nuclease digestion, unlike untreated cells or HU-treated wild-type MEFs. Interestingly, *Brca2^+/Δ27^* and *Brca2^+/SA^* MEFs also showed reduced CldU/IdU ratios (**Figures 2A, S2C and S2D**). Thus, both homozygous and heterozygous mutation of the BRCA2 C-terminal RAD51 interaction site results in the accumulation of ssDNA gaps.

**Figure 2.**
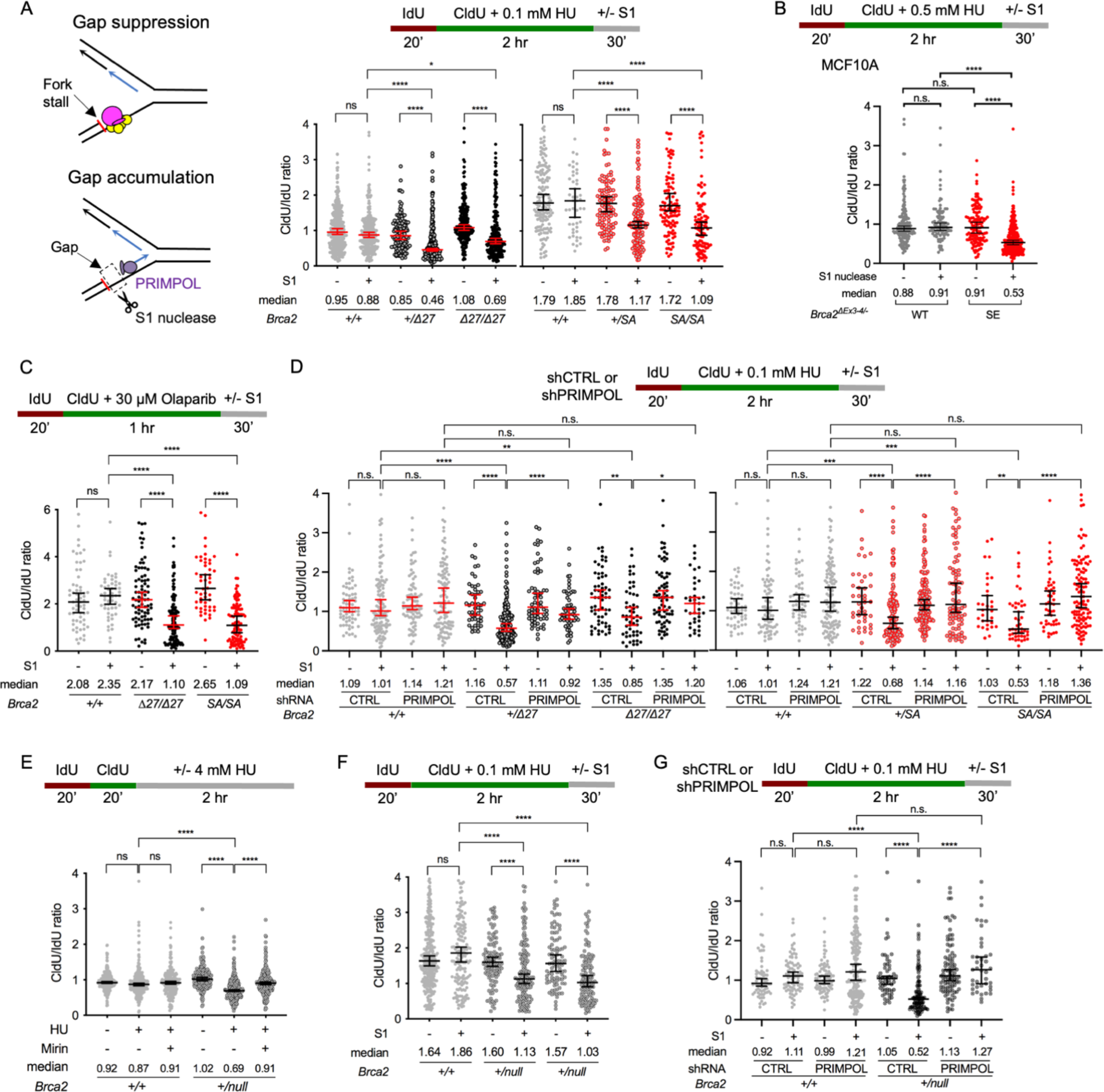
BRCA2 C terminal RAD51 interaction site suppresses PRIMPOL-mediated ssDNA gaps. **A.** Impaired gap suppression in *Brca2* homozygous and heterozygous cells treated with HU. Diagram shows PRIMPOL activity creating S1 nuclease-sensitive ssDNA gaps when BRCA2 and RAD51 are absent. Primary MEFs were pulsed with IdU for 20 min and then CldU together with 0.1 mM HU for 2 hr. Cells were harvested and treated with S1 nuclease for 30 min to digest DNA fibers with ssDNA gaps. Statistical analysis, Mann- Whitney U test (median ± 95% confidence intervals). ns, no significance; *, p<0.05; **, p<0.01; ****, p<0.0001. **B.** Human mammary MCF10A cells overexpressing BRCA2 S3291E show impaired gap suppression. DNA fiber assays were similar to (**A**) except with 0.5 mM HU. **C.** Impaired gap suppression in *Brca2^Δ27/Δ27^* and *Brca2^SA/SA^* cells treated with olaparib. DNA fiber assays were similar to (**A**) but with 30 µM olaparib treatment instead of HU. Results are combined from two experiments. **D.** PRIMPOL knockdown suppresses gap formation in *Brca2* mutant cells. DNA fiber assays were similar to (**A**) but with immortalized MEFS stably expressing shRNAs for *Primpol* knockdown or control (CTRL). Statistical analysis, Mann-Whitney U test (median ± 95% confidence intervals). ns, not significant; *, p<0.05; **, p<0.01; ***, p<0.001; ****, p<0.0001. **E.** Stalled fork degradation in *Brca2^+/null^* primary MEFs. DNA fiber assays were similar to Figure 1D. **F.** Impaired gap suppression in *Brca2^+/null^* primary MEFs. DNA fiber assays were similar to (**A**). **G.** PRIMPOL knockdown suppresses gap formation in *Brca2^+/null^* immortalized MEFs. DNA fiber assays were similar to (**A**). See **Figure S2C,D,F,H-K** for replicates.

To ascertain if the observations we made in murine cells are pertinent to human cells, we probed stalled fork protection and gap suppression in MCF10A cells, a non-transformed human mammary epithelial cell line. We used BRCA2 mutant MCF10A cells complemented with BRCA2 S3291E (SE), which fails to rescue fork protection but suppresses replication stress (Feng and Jasin, 2017) (**Figure S2E**). As with the *Brca2* mutant MEFs, we found that these cells accumulated ssDNA gaps upon exposure to HU (**Figures 2B** **and** **S2F**). These results suggest that loss of RAD51 interaction at the BRCA2 C terminus leads to fork protection and gap suppression defects in both murine and human cells.

BRCA-deficient cells accumulate gaps in response to the poly(ADP-ribose) polymerase inhibitor (PARPi) olaparib (Cong *et al*., 2021). To determine whether the BRCA2 C-terminal RAD51 interaction site is essential to prevent PARPi-induced gap accumulation, we repeated our experiments with olaparib (**Figure 2C**). Similar to HU treatment, both *Brca2^Δ27/Δ27^* and *Brca2^SA/SA^* MEFs showed reduced CldU/IdU ratios upon S1 nuclease digestion, unlike untreated cells or olaparib-treated wild-type MEFs, indicating that ssDNA gap accumulation is a general outcome of replication stress when the BRCA2 C terminus is compromised.

In the absence of BRCA2, PRIMPOL repriming bypasses DNA lesions, leading to ssDNA gap formation (Taglialatela *et al*., 2021; Tirman *et al*., 2021). Thus, we downregulated PRIMPOL expression in immortalized MEFs and measured S1 nuclease-sensitive DNA tracts upon HU treatment (**Figures 2D**, **S2G** **and** **S2H**). Similar to primary MEFs, S1 nuclease-digested immortalized homozygous and heterozygous *Brca2* mutant MEFs showed a reduced CldU/IdU ratio compared to undigested counterparts and S1 nuclease-digested wild-type cells. Importantly, with PRIMPOL depletion, CldU/IdU tract ratios were largely restored to normal levels. PRIMPOL knockdown also restored CldU/IdU ratios in the presence of olaparib (**Figure S2I**). Collectively, our results suggest that BRCA2 C-terminal RAD51 interaction is crucial for preventing PRIMPOL-mediated ssDNA gap accumulation.

### Haploinsufficiency for BRCA2-mediated stalled fork protection and gap suppression

In *Brca2* heterozygous cells, defective replication fork protection and gap suppression may arise from either haploinsufficiency of the wild-type *Brca2* copy or dominant negative effects of the mutant alleles. To evaluate these possibilities, we investigated cells containing a *Brca2* null allele (Ludwig *et al*., 2001). In fork protection assays, we found that HU-treated *Brca2^+/null^* primary MEFs had a significantly reduced CldU/IdU tract length ratio in comparison to untreated cells and HU-treated wild-type MEFs (**Figures 2E** **and** **S2J**), similar to *Brca2^+/SA^* and *Brca2^+/Δ27^* cells. This defect in replication fork protection is rescued by mirin treatment. Likewise, in *Brca2^+/null^* primary MEFs, we saw a reduction in the CldU/IdU ratio with S1 nuclease digestion compared to wild-type MEFs treated with S1 nuclease (**Figures 2F** **and** **S2K**). Knocking down PRIMPOL in immortalized MEFs restored the CldU/IdU ratio to normal levels (**Figures 2G** **and** **S2H**). These results indicate that *Brca2* haploinsufficiency causes both defective replication fork protection and ssDNA gap suppression and thus may be sufficient to explain the defects seen in heterozygous *Δ27* and *SA* cells.

### Mild genomic instability in HU-treated MEFs with fork protection and gap suppression defects

To ascertain which function(s) of BRCA2 are required to suppress DNA damage during unperturbed or replication stress conditions, we evaluated the frequency of chromosome aberrations in metaphase spreads of untreated and HU-treated primary MEFs (**Figures 3A** **and** **3B**). We found that each of the HDR-proficient but fork protection and gap suppression-deficient MEFs – *Brca2^+/Δ27^*, *Brca2^+/SA^*, *Brca2^SA/SA^* and *Brca2^+/null^*– showed a small but statistically significant increase in spontaneous and HU-induced chromosome aberrations (mean 0.8 and 1.6 aberrations per 40 chromosomes, respectively). However, substantially higher numbers of aberrations were observed in the *Brca2^Δ27/Δ27^* MEFs (mean 2.2 and 4.0 aberrations, respectively), which are deficient in HDR, fork protection, and gap suppression. Most of the HU-induced damage incurred in all mutants comprised chromosome breaks and gaps and acentric chromosome fragments (**Figure 3B**). In addition, *Brca2^Δ27/Δ27^* MEFs accumulated chromosome exchanges and radials. Thus, these results suggest that fork protection and gap suppression play much less of a role in suppressing chromosome damage relative to HDR.

**Figure 3.**
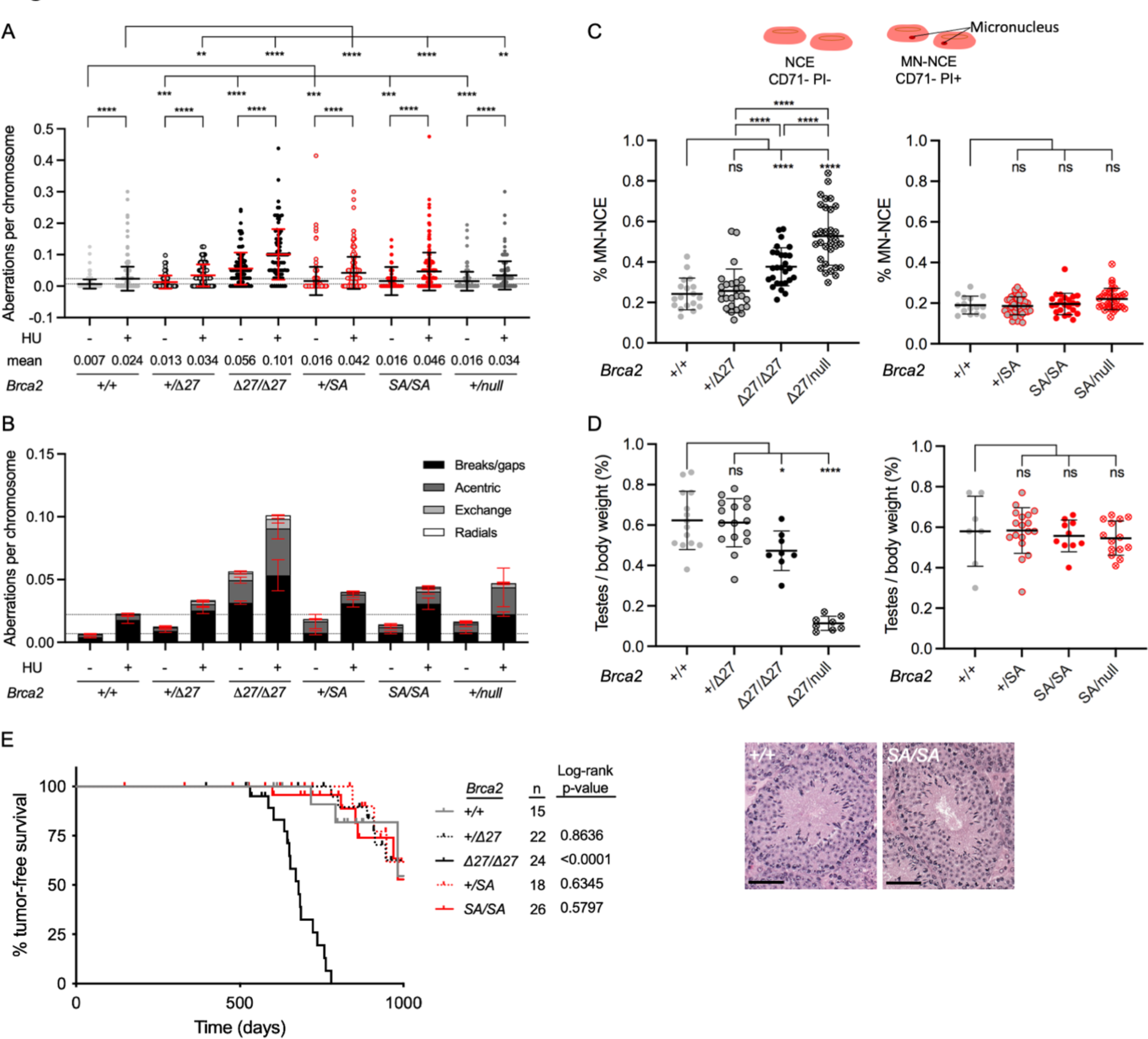
BRCA2 C-terminal RAD51 interaction site is dispensable for maintaining genome integrity. **A.** Spontaneous and HU-induced chromosome aberrations are substantially increased in HDR-deficient *Brca2^Δ27/Δ27^*primary MEFs but only mildly in *Brca2* mutants proficient in HDR. Three repeats were performed for each genotype. Statistical analysis, unpaired Student’s t test (mean ± 1 SD). ns, not significant; *, p<0.05, **, p<0.01, ***, p<0.001, ****, p<0.0001. **B.** Breakdown of chromosomal aberrations in (**A**). Most are breaks/gaps, although acentrics, exchanges, and radials are also observed, especially in *Brca2^Δ27/Δ27^*MEFs. **C.** Micronuclei are observed in *Brca2^Δ27/Δ27^* and especially *Brca2^Δ27/null^* blood cells, but not in *Brca2^SA/SA^* or *Brca2^SA/SA^* blood cells. Normochromic erythrocytes (NCEs) are negative for both CD71 and propidium iodide (PI) but become PI positive when a micronucleus is present (MN-NCE). (See also **Figure S3B**.) Peripheral blood was collected from 1.5 to 4-month old mice and stained for CD71 and PI. MN-NCE by sex is in Figure S3C,D.) Statistical analysis, unpaired Student’s t test (mean ± 1 SD). ns, not significant; ****, p<0.0001. **D.** Testis weights are reduced in *Brca2^Δ27/Δ27^* and especially *Brca2^Δ27/null^* mice, but not in *Brca2^SA/SA^* or *Brca2^SA/SA^* mice. For each mouse (2- to 4-month-old), the combined weight from both testes is presented as a percentage of body weight. (See body weights and testis weights in **Figure S3A,E**.) Statistical analysis, unpaired Student’s t test (mean ± 1 SD). ns, not significance; *, p<0.05; ****, p<0.0001. Inset images are representative stage IV cross sections of H&E-stained tubules from *Brca2^+/+^* and *Brca2^SA/SA^*testes. Additional genotypes are in **Figure S3F**. Scale bar, 100 µm. **E.** *Brca2^Δ27/Δ27^* mice, but not other *Brca2* genotypes, are tumor prone. Kaplan-Meier tumor-free survival curves were analyzed using the log-rank (Mantel-Cox) test.

### HDR-proficient, but fork protection- and gap suppression-deficient, mice are genetically stable

We next investigated the effects of fork protection and gap suppression deficiencies on mouse phenotypes. No discernible developmental issues or physical deformities were observed in *Brca2^SA/SA^* mice and they also had a normal body size, as was also the case for the other *Brca2* genotypes, including *Brca2^Δ27/Δ27^* mice (**Figure S3A**).

General genomic instability in the mice was measured by assessing micronucleus formation. Micronucleated, normochromic erythrocytes (MN-NCE) can be detected by staining peripheral blood for CD71 and propidium iodide (Shima *et al*., 2003). Mice with fork protection and gap suppression defects – *Brca2^+/Δ27^*, *Brca2^+/SA^*, *Brca2^SA/SA^*and *Brca2^SA/null^* – showed normal levels of MN-NCEs (**Figures 3C** **and** **S3B**). This is consistent with the low frequency of chromosomal aberrations seen in MEFs, especially considering that acentric fragments are more likely to be found in micronuclei (Shimizu *et al*., 1996) (**Figure 3B**). By comparison, *Brca2^Δ27/Δ27^* mice had a pronounced increase in MN-NCEs (1.5-fold), which was further exacerbated in *Brca2^Δ27/null^* mice (2.2-fold). The increase was observed in both females and males, the latter of which have a higher baseline level of MN-NCEs (**Figures S3C** **and** **S3D**). Taken together, these results reinforce the importance of HDR for maintaining genomic stability.

Due to its key role in HDR, BRCA2 is required for meiotic progression in mice (Sharan *et al*., 2004). While *Brca2^Δ27/Δ27^* mice are fertile, they have a 24% reduced average testis weight as compared with wild-type or heterozygous mice (Abreu *et al*., 2018) (**Figures 3D** **and** **S3E**). We found this to be greatly exacerbated in *Brca2^Δ27/null^*mice, which had a 6-fold reduction in testis weight (**Figures 3D** **and** **S3E**). By contrast, *Brca2^SA/SA^* mice had normal testis weights, as did *Brca2^SA/null^*mice, indicating that the *SA* allele is not sensitive to a reduced dosage. Seminiferous tubules of *Brca2^SA/SA^*, *Brca2^SA/null^*, and *Brca2^Δ27/27^* mice progressed through various stages of spermatogenesis, and the epididymis of each showed abundant sperm (F**igure 3D** **inset;** **Figure S3F**). However, the seminiferous tubules of *Brca2^Δ27/null^*mice were missing or reduced for many germ cell stages and the epididymis was devoid of sperm, likely due to both spermatogonial and spermatocyte defects. Thus, unlike HDR defects, fork protection and gap suppression defects do not compromise the proliferation or differentiation of germ cells in male mice.

### Spontaneous tumor formation is accelerated only in *Brca2^Δ27/Δ27^* mice

*Brca2^Δ27/Δ27^* mice have been reported to be tumor prone (McAllister *et al*., 2002). However, it is unclear whether defects in replication fork protection and gap suppression predispose to tumorigenesis. To address this, we monitored cohorts of mice with *Δ27* and *SA* hetero- and homozygosity for tumor formation (**Figure 3E**). As reported, *Brca2^Δ27/Δ27^*mice had significantly shorter tumor latency compared to wild-type littermates (679 day median survival versus >1000 days, respectively). However, none of the *Brca2^SA/SA^*, *Brca2^+/SA^*, or *Brca2^+/Δ27^*mice was significantly different from their wild-type littermates. Thus, BRCA2-mediated replication fork protection and ssDNA gap suppression are not critical for preventing tumorigenesis, while an associated HDR defect is tumor predisposing.

### Loss of BRCA2-mediated HDR substantially increases sensitivity to olaparib and cisplatin

We next examined whether HU-induced nascent strand degradation and ssDNA gap accumulation is sufficient to cause increased HU sensitivity. Immortalized MEFs were treated with increasing concentrations of HU for 24 hr, and the number of viable cells was measured after 6 days. *Brca2^Δ27/Δ27^*MEFs showed a mild sensitivity to HU treatment compared to wild-type cells (LD50s: *Brca2^Δ27/Δ27^*, 81.7 ± 4.0 µM versus *Brca2^+/+^*, 118.4 ± 12.6 µM; two-way ANOVA, p<0.0001) (**Figures 4A** **and** **S4A**). By contrast, *Brca2^+/Δ27^* and *Brca2^+/SA^*MEFs showed similar HU sensitivity as wild type, while *Brca2^SA/SA^*MEFs were slightly more resistant (**Figures 4A** **and** **S4A**). Thus, even though HU leads to stalled fork degradation and gap accumulation with an associated low level of chromosome aberrations in *Brca2^+/Δ27^*, *Brca2^+/SA^*, and *Brca2^SA/SA^* MEFs, these cells do not show heightened sensitivity to HU.

**Figure 4.**
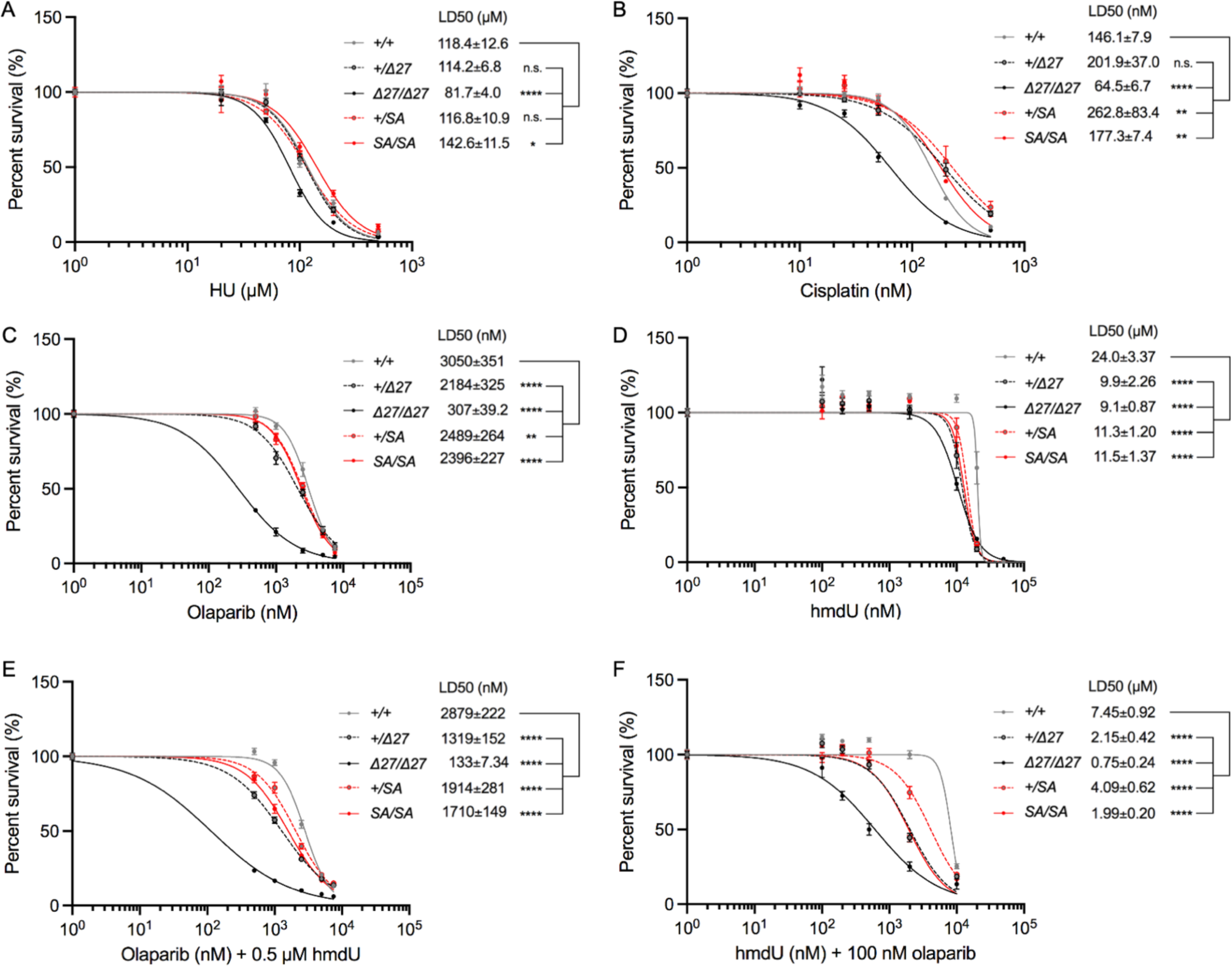
Genotoxin sensitivity is greatest in the absence of HDR, but is observed for hmdU in fork protection/gap suppression defective cells. **A-F.** Genotoxin sensitivity assays of immortalized MEFs. Cells were treated with various concentrations of HU (**A**), cisplatin (**B**), olaparib (**C**), hmdU (**D**), olaparib with a fixed concentration of hmdU (**E**), or hmdU with a fixed concentration of olaparib (**F**) for 6 days, and their survival was plotted. In all conditions, HDR-deficient *Brca2^Δ27/Δ27^* MEFs display the greatest sensitivity, while MEFs with only fork protection/gap suppression defects show milder or no sensitivity, except in the case of hmdU alone, in which the sensitivities of all *Brca2* mutants are more similar (**D**) and are exacerbated by olaparib (**F**). Statistical analysis, two-way ANOVA test (mean ± SEM). ns, not significant; *, p<0.05; **, p<0.01; ***, p<0.001; ****, p<0.0001.

To determine if this lack of sensitivity is applicable to other agents that cause replication stress, we treated the same set of immortalized MEFs with cisplatin, which leads to DNA crosslinks, in this case continuously over 6 days. As expected, *Brca2^Δ27/Δ27^*MEFs were significantly more sensitive to cisplatin treatment than wild type (LD50s: *Brca2^Δ27/Δ27^*, 64.50 ± 6.7 nM versus *Brca2^+/+^*, 146.1 ± 7.9 nM; two-way ANOVA, p<0.0001) (**Figures 4B** **and** **S4B**). However, as with HU, the fork protection and gap suppression mutants were not more sensitive (**Figures 4B** **and** **S4B**).

BRCA2-deficient cells are exquisitely sensitive to inhibitors which trap PARP on DNA (Bryant *et al*., 2005; Farmer *et al*., 2005; Murai *et al*., 2012). This sensitivity was initially attributed to loss of HDR, due to an inability to repair collapsed replication forks, but recent investigations have suggested that fork protection and ssDNA gap suppression may play more prominent roles (Cong *et al*., 2021; Ray Chaudhuri *et al*., 2016). To test this in an isogenic series, we treated immortalized MEFs continuously with olaparib for 6 days. While all fork protection and gap suppression mutant MEFs showed some sensitivity to olaparib compared to wild type, only the additionally HDR-deficient *Brca2^Δ27/Δ27^* cells showed substantial sensitivity (**Figure 4C**). Specifically, *Brca2^Δ27/Δ27^* MEFs were about 10-fold more sensitive than wild type when comparing LD50s (307.5 ± 39.2 nM versus 3050 ± 351 nM, respectively), whereas *Brca2^SA/SA^*, *Brca2^+/SA^*, and *Brca2^+/Δ27^* MEFs were only 1.2- to 1.4- fold more sensitive (2396 ± 227 nM, 2489 ± 264 nM, and 2184 ± 325 nM, respectively) (**Figures 4C** **and** **S4C**). Thus, when PARP is inhibited, HDR appears to have a major role in mitigating cell death, while fork protection and gap suppression play more minor roles.

### Loss of BRCA2-mediated fork protection/gap suppression leads to hmdU sensitivity

In addition to PARP inhibitors, other agents are being developed to specifically target BRCA1 and BRCA2 mutant cells, including the cytotoxic nucleotide 5-hydroxymethyl-2’-deoxyuridine (hmdU) (Fugger *et al*., 2021). The introduction of hmdU into DNA in replicating cells, through disruption of the nucleotide salvage pathway and/or direct addition to the nucleotide pool, has recently been shown to sensitize BRCA1/2-deficient cells, especially in conjunction with PARP inhibition. To determine which function of BRCA2 is important in this context, we tested the effect of hmdU on our separation-of-function MEFs. Surprisingly, all *Brca2* mutant MEFs were more sensitive to hmdU than wild-type MEFs, with 2.1- to 2.6-fold lower LD50s (wild type, 24.0 ± 3.37 µM; *Brca2^SA/SA^*, 11.5 ± 1.37 µM; *Brca2^+/SA^*, 11.3 ± 1.20 µM; *Brca2^Δ27/Δ27^*, 9.1 ± 0.87 µM; *Brca2^+/Δ27^*, 9.9 ± 2.26 µM) (**Figures 4D** **and** **S4D**). This result suggests that BRCA2-mediated fork protection and/or gap suppression dictate sensitivity to hmdU.

Abasic sites generated at hmdU have been predicted to lead to PARP trapping (Fugger *et al*., 2021). To determine whether hmdU can enhance sensitivity to PARP inhibition, we treated MEFs with increasing doses of olaparib in the presence of a non-lethal dose of hmdU (0.5 µM) when administered on its own (**Figures 4E** **and** **S4G**). Interestingly, all *Brca2* mutant MEFs became more sensitive to olaparib plus hmdU (**Figures 4E** **and** **S4E**). *Brca2^Δ27/Δ27^* cells were by far the most sensitive, with a 21-fold greater sensitivity than wild type (LD50s: 132.8 ± 7.34 nM versus 2879 ± 222 nM, respectively; **Figure S4E**). *Brca2^+/Δ27^*, *Brca2^+/SA^* and *Brca2^SA/SA^* MEFs also became more sensitive, although to a lesser degree, by about 1.5- to 2.2-fold (1319 ± 152 nM, 1914 b44 b4± 281 nM and 1710 ± 149 nM, respectively).

Conversely, we also treated cells with increasing doses of hmdU in the presence of a low dose of olaparib (100 nM). All cell lines showed increased sensitivity to hmdU in the presence of this dose of olaparib (**Figures 4F** **and** **S4G**). While *Brca2^Δ27/Δ27^* cells were the most sensitive, *Brca2^SA/SA^*, *Brca2^+/SA^*, and *Brca2^+/Δ27^*, MEFs were also significantly more sensitive than wild type. *Brca2^Δ27/Δ27^*cells were about 10-fold more sensitive to hmdU plus 100 nM olaparib (LD50s: 0.75 ± 0.24 µM versus 7.45 ± 0.92 µM, respectively), whereas *Brca2^SA/SA^*, *Brca2^+/SA^*, and *Brca2^+/Δ27^*, cells were 1.8- to 3.5-fold more sensitive (1.99 ± 0.20 µM, 4.09 ± 0.62 µM, and 2.15 ± 0.42 µM, respectively; **Figure S4F**). Thus, *Brca2* mutants defective in fork protection and ssDNA gap suppression can be targeted with combination therapy of a PARP inhibitor and hmdU.

### Defects in gap suppression dictate sensitivity to olaparib and hmdU, but only when HDR is proficient

As sensitivity to combined olaparib/hmdU treatment is linked to mutants that are defective in both fork protection and gap suppression, we sought to establish which pathway defect may be more important for drug sensitivity. To that end, we separated these two pathways by perturbing genes that regulate them. PRIMPOL knockdown prevents gap accumulation in our system (**Figures 2C**, **2G** **and** **S2F**). To determine if stalled fork protection is also impacted by PRIMPOL knockdown, we performed DNA fiber assays after HU treatment and found that CldU/IdU ratios were still reduced in *Brca2^Δ27/Δ27^*, *Brca2^SA/SA^*, and *Brca2^+/null^* immortalized MEFs (**Figure 5A**). Thus, stalled fork degradation still occurs with PRIMPOL knockdown, consistent with what has been previously been reported for BRCA1-deficient cells (Quinet *et al*., 2020).

**Figure 5.**
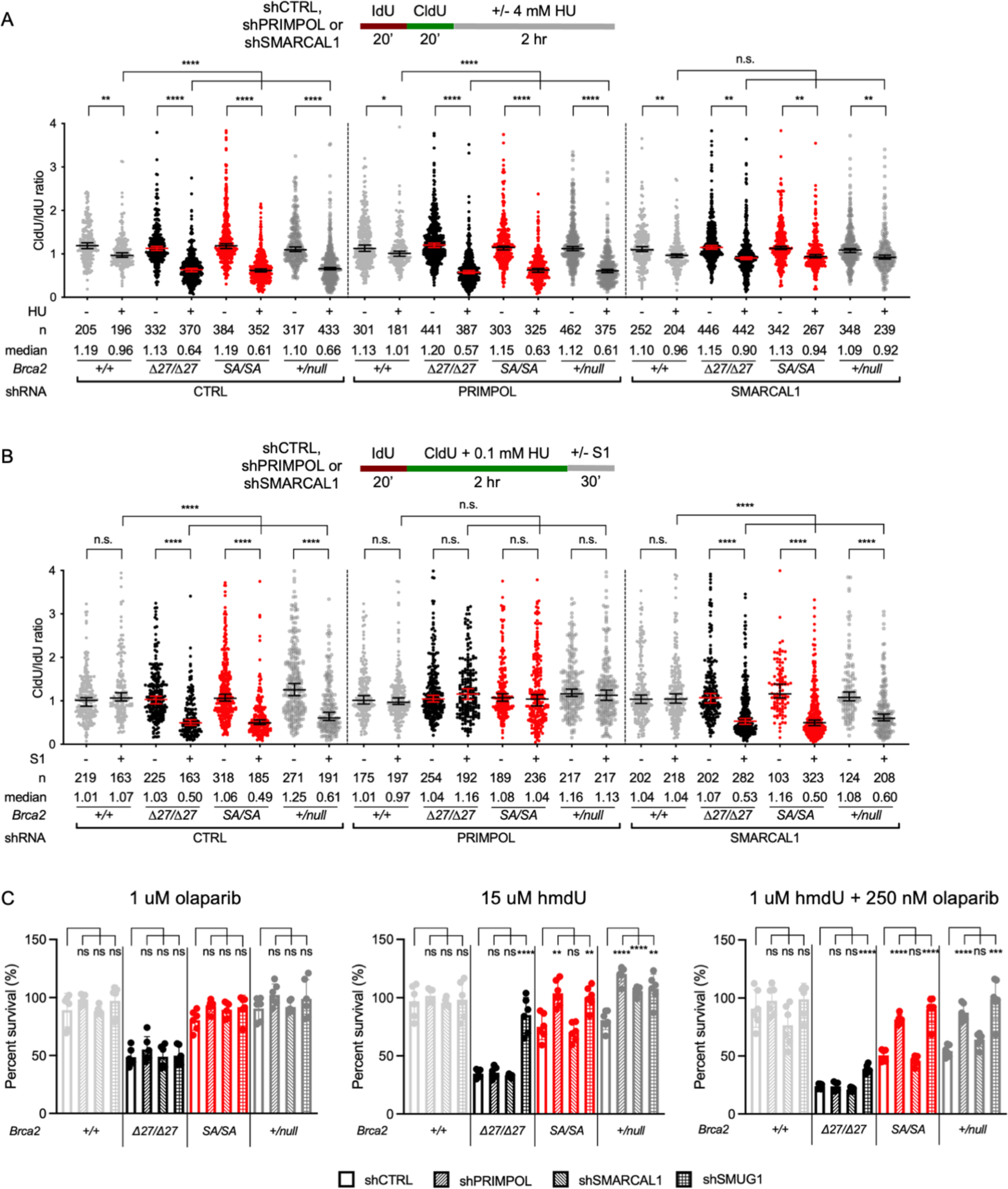
hmdU sensitivity arises from PRIMPOL activity in HDR-proficient cells, but not in HDR- deficient cells. **A.** Stalled fork protection is restored in *Brca2* mutants with SMARCAL1, but not PRIMPOL, knockdown. DNA fiber assays were performed in immortalized MEFs stably expressing the indicated shRNAs. Statistical analysis, Mann-Whitney U test (median ± 95% confidence intervals). ns, not significant; *, p<0.05; **, p<0.01; ****, p<0.0001. **B.** Gap suppression is restored in *Brca2* mutants with PRIMPOL, but not SMARCAL1, knockdown. DNA fiber assays were performed in immortalized MEFs stably expressing the indicated shRNAs. Statistical analysis, Mann-Whitney U test (median ± 95% confidence intervals). ns, not significant; ****, p<0.0001. **C.** hmdU sensitivity arises from PRIMPOL activity in HDR-proficient cells, but not in HDR-deficient cells. Olaparib sensitivity is not affected by any of the knockdowns. However, hmdU sensitivity is rescued by SMUG1 knockdown in all mutants, indicating that sensitivity arises from abasic sites formed at hmdU incorporated in the genome. Moreover, gaps arising from PRIMPOL activity are responsible for the hmdU sensitivity of *Brca2^SA/SA^* and *Brca2^+/null^* cells, but not HDR-deficient *Brca2^Δ27/Δ27^* cells. Statistical analysis, unpaired Student’s t test (mean ± 1 SD). ns, not significance; *, p<0.05; ***, p<0.001; ****, p<0.0001.

Conversely, the motor protein SMARCAL1 participates in the reversal of stalled replication forks, which provides the substrates for degradation by the MRE11 nuclease (Kolinjivadi *et al*., 2017). Thus, SMARCAL1 knockdown rescues stalled fork protection in BRCA1/2-deficient cells (Taglialatela *et al*., 2017). We verified that SMARCAL1 knockdown prevented fork degradation in *Brca2^Δ27/Δ27^*, *Brca2^SA/SA^* and *Brca2^+/null^* immortalized MEFs (**Figures 5A** **and** **S4H**). However, S1-sensitive gaps were still observed, indicating that SMARCAL1 does not affect the gap suppression defect in these cells (**Figure 5B**).

The uracil glycosylase SMUG1 hydrolyzes hmdU to create abasic sites, which are considered to be the toxic lesion because they can lead to replication fork collapse (Fugger *et al*., 2021). We depleted SMUG1 in each of the mutants and confirmed that abasic sites are the source of their sensitivity to hmdU, but as expected it did not affect olaparib sensitivity (**Figure 5C**).

Having separated the fork protection and gap suppression defects, as well as determining the cause hmdU sensitivity, we asked how each pathway impacts olaparib and/or hmdU sensitivities. Strikingly, the strong sensitivity of *Brca2^Δ27/Δ27^* MEFs to either drug alone or combined was not affected by PRIMPOL or SMARCAL1 knockdown, implying that reviving fork protection or gap suppression is not enough to rescue chemosensitivity when HDR is deficient (**Figure 5C**). Conversely, in *Brca2^SA/SA^* and *Brca2^+/null^* MEFs, knocking down PRIMPOL fully rescued their sensitivity to hmdU, and partially to combination treatment, but not to olaparib alone. This suggests that when HDR is proficient, PRIMPOL-mediated ssDNA gap accumulation at abasic sites causes drug sensitivity. However, preventing fork degradation with SMARCAL1 knockdown for the most part had no effect. We did notice a slight rescue of *Brca2^+/null^* MEF sensitivity to hmdU with SMARCAL1 knockdown, but not to the combined treatment, suggesting an hmdU-specific defect. Thus, HDR has a dominant role in suppressing sensitivity to olaparib and hmdU. However, gap suppression promotes hmdU resistance when HDR is intact.

## Discussion

### BRCA2 RAD51-interacting carboxyl-terminal domain mediates stalled fork protection and ssDNA gap suppression functions

The role of BRCA2 in maintaining genome integrity is integral to the suppression of tumorigenesis. While its role in HDR was established more than two decades ago (Moynahan *et al*., 2001), recent work has called into question the importance of BRCA2 function in HDR, given the participation of BRCA2 in other genome maintenance pathways (Cong and Cantor, 2022). These other pathways involve the protection of stalled replication forks from MRE11 nuclease degradation (Schlacher *et al*., 2011) and the suppression of ssDNA gap formation (Panzarino *et al*., 2021; Taglialatela *et al*., 2021; Tirman *et al*., 2021). BRCA2 either prevents or protects gaps on the leading strand formed by PRIMPOL-mediated repriming of DNA synthesis downstream of stalled replication forks (Kang *et al*., 2021; Tirman *et al*., 2021). BRCA2 has also been proposed to prevent lagging strand gaps at sites of Okazaki fragment processing (Cong *et al*., 2021), but such gaps were not detected in our system, given that PRIMPOL is responsible for the S1 nuclease sensitive gaps we observed. Given that genome instability and therapy response could arise due to abrogation of any one of these pathways, we set out to determine the relative contribution of these pathways.

To that end, we generated a BRCA2 separation-of-function mutation in mouse that is functional for HDR but leads to defective fork protection (S3214A or “SA”) (**Figure 6**). This mutation, which is based on our previous findings in human and hamster cells (Esashi *et al*., 2005; Feng and Jasin, 2017; Schlacher *et al*., 2011), disrupts RAD51 interaction at the C terminus but does not affect RAD51 interaction sites – the BRC repeats – in the central region of BRCA2 that are critical for HDR. Unexpectedly, we discovered that mutation of this site also leads to the accumulation of ssDNA gaps due to PRIMPOL activity during replication stress (**Figure 6**). Gaps were also observed in human mammary cells expressing a similar separation-of-function mutation at the cognate BRCA2 residue (SE), which we previously reported was able to promote HDR and suppress replication stress (Feng and Jasin, 2017).

**Figure 6.**
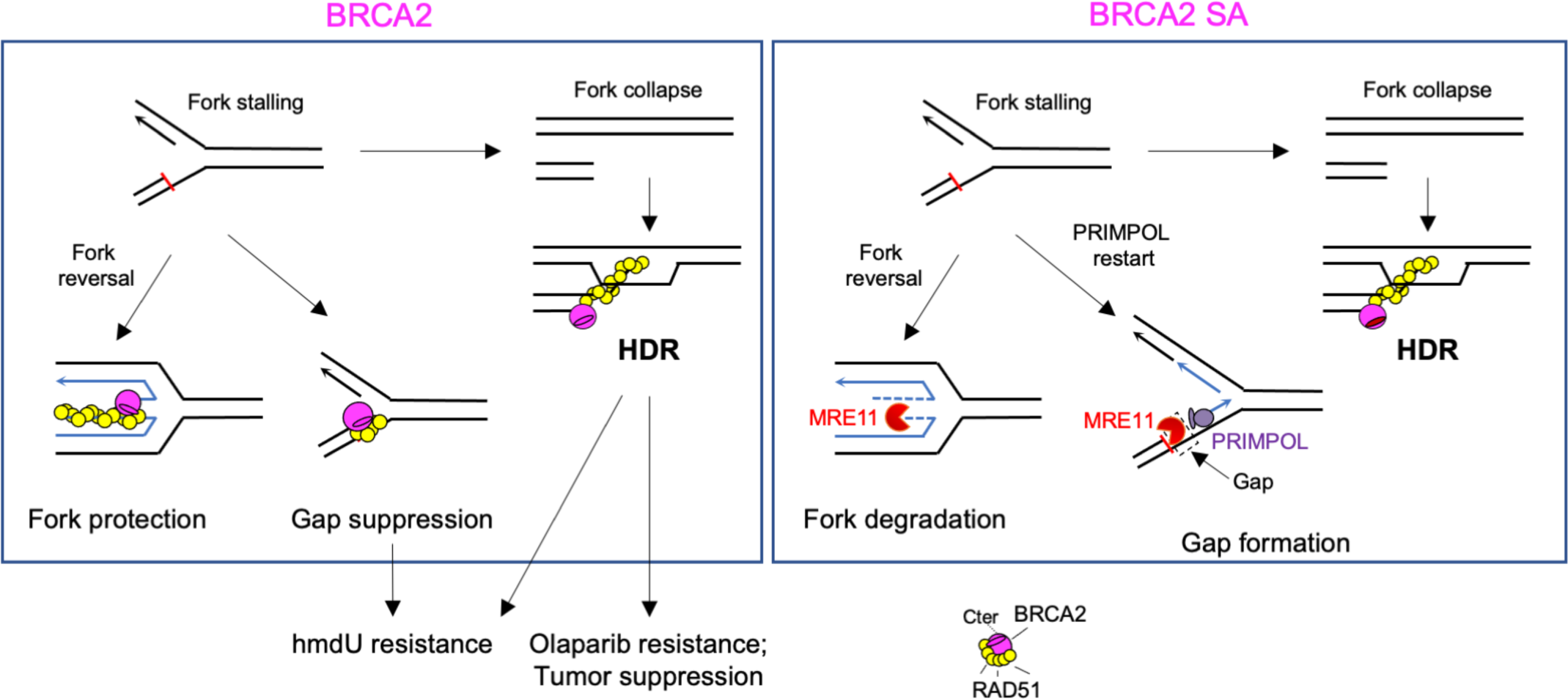
BRCA2 function during replication stress. HDR is required when replication forks collapse, including with olaparib treatment. However, when forks stall, BRCA2 protects reversed forks, which can restart when the replication stress is relieved, and prevents or protects PRIMPOL-mediated ssDNA gaps. Loss of the BRCA2 C-terminal RAD51-interacting site leads to fork degradation and ssDNA gap formation. Gap suppression and HDR are important for hmdU resistance, while HDR plays the major role in olaparib resistance and tumor suppression.

RAD51 interaction at the BRCA2 C terminus stabilizes RAD51 filaments on ssDNA and dsDNA (Davies and Pellegrini, 2007; Esashi *et al*., 2007; Halder *et al*., 2022), while the BRC repeats are important for nucleation of RAD51 nucleoprotein filaments. The requirement for an intact C-terminal site for fork protection has been explained by BRCA2-mediated RAD51 filament stabilization (Schlacher *et al*., 2011), specifically at reversed forks (Mijic *et al*., 2017), to block MRE11-mediated nucleolytic degradation. RAD51 filament stabilization specifically on dsDNA may be particularly important during fork protection (Halder *et al*., 2022). Perhaps BRCA2 similarly stabilizes RAD51 at dsDNA adjacent to gaps to prevent MRE11 nuclease activity and to assist downstream processes to fill the gaps (Tirman *et al*., 2021) (**Figure 6**).

In contrast to the SA point mutation, deletion of the C terminus (Δ27) causes HDR defects (Kass *et al*., 2016; Moynahan *et al*., 2001), in addition to fork protection and gap suppression defects. The Δ27 deletion includes a KKRR motif which has recently been reported to bind DNA and promote HDR as well as fork protection (Kwon *et al*., 2023). This motif is intact in the SA mutant. Interestingly, loss of this motif together with the RAD51 interaction site in the Δ27 mutant causes similarly severe defects in fork protection and gap suppression as SA mutation.

### BRCA2 heterozygosity results in defects in stalled fork protection and gap suppression

Recent reports have emphasized the importance of normal levels of BRCA2 protein in genome maintenance. Aldehyde-induced BRCA2 degradation in BRCA2 heterozygous cells leads to loss of fork protection not seen in wild-type cells (Tan *et al*., 2017). Moreover, sequencing data from breast tissue of BRCA2 mutation carriers has shown elevated sub-chromosomal copy number variations not seen in non-carriers (Karaayvaz-Yildirim *et al*., 2020). In the course of our studies, we discovered that BRCA2 heterozygosity drives deficiencies in both fork protection and gap suppression without affecting HDR. All three BRCA2 heterozygous mutants are defective in these two pathways, with similar levels of deficiencies in the homozygous and heterozygous mutants. Moreover, as these defects are observed when one allele of BRCA2 is null, haploinsufficiency is implied.

The requirement for normal levels of wild-type BRCA2 can be interpreted as BRCA2 having a stoichiometric role in fork protection/gap suppression, whereas during HDR, lower BRCA2 levels may be sufficient to promote the catalytic function of RAD51. Such a protective role for BRCA2 in conjunction with RAD51 is consistent with the finding that over-expression of wild-type RAD51 or a non-enzymatic RAD51 mutant that can still form filaments protects stalled forks (Mason *et al*., 2019; Schlacher *et al*., 2012). Further, biochemical studies have demonstrated that RAD51 filaments stabilized on dsDNA by stoichiometric amounts of the BRCA2 C-terminal peptide protect against MRE11 nuclease activity when in competition with a RAD51 filament destabilizing BRC4 peptide (Halder *et al*., 2022).

### Loss of BRCA2-mediated stalled fork protection and gap suppression does not lead to genomic instability or accelerated tumor formation in mice

When compared to the BRCA2 Δ27 deletion, the SA mutation causes minimal genome instability, as evidenced by no increase in micronuclei in peripheral blood, minimal increase in chromosomal aberrations in MEFs, normal testis weights, and unremarkable sensitivity to conventional genotoxic agents. All three heterozygous mutants show similar phenotypes where tested. Moreover, while tumor latency for *Brca2^Δ27/Δ27^* mice is significantly shorter than for wild-type mice, it is not affected in the fork protection/gap suppression mutants with intact HDR. Therefore, we conclude that of the three functions, HDR is the predominant BRCA2-mediated function involved in genome maintenance and tumor suppression.

Our findings are congruent with studies of separation-of-function alleles in BRCA1-BARD1 mutant mice and non-transformed BRCA2 mutant human mammary epithelial cells, both of which have reported a greater reliance on HDR over fork protection for genome integrity and cell viability, as well as for tumor suppression in the case of the BRCA1-BARD1 mutant mice (Billing *et al*., 2018; Feng and Jasin, 2017). However, restoration of fork protection in *Brca1* and *Brca2*-deficient mouse B cells via *Parp1*, *Ptip* or *Chd4* knockout has been reported to be sufficient to suppress olaparib and cisplatin-induced chromosome aberrations without restoration of HDR, suggesting that fork protection can play an important role in maintaining genome integrity in some contexts (Ding *et al*., 2016; Ray Chaudhuri *et al*., 2016). Complicating matters further, knockdown of PRIMPOL in BRCA1 and BRCA2 deficient human tumor cells has been reported to cause higher genome instability and decreased cell viability, suggesting that certain tumor cell lines *actually* require PRIMPOL activity to survive in the absence of HDR (Kang *et al*., 2021; Quinet *et al*., 2020; Taglialatela *et al*., 2021). These inconsistencies may be attributable to different biological contexts in the studies. Nevertheless, it is clear from the work shown here that, under normal physiological conditions in mice, loss of both fork protection and gap suppression does not cause profound genomic instability or predispose to tumorigenesis, unlike HDR perturbation.

### PRIMPOL-dependent ssDNA gaps and sensitivity to hmdU

The correlation between BRCA2 functionality and response to chemotherapy is clinically relevant. In this study, we found that fork protection/gap suppression defective *Brca2* mutant cells are sensitive to exogenous treatment with the cytotoxic nucleotide hmdU. Sensitivity to increased levels of genomic hmdU by HDR-deficient cells was recently uncovered in a synthetic lethal screen (Fugger *et al*., 2021). Endogenously, hmdU accumulates in genomic DNA through several ways, including loss of the glycosidase DNPH1, which hydrolyzes hmdU, hydroxylation of thymidine by TET dioxygenases, oxidation of thymidine by reactive oxygen species, and deamination of 5-hydroxymethylcytosine (5hmC) by cytidine deaminase (Fugger *et al*., 2021; Olinski *et al*., 2016; Pfaffeneder *et al*., 2014; Teebor *et al*., 1984). Although not cytotoxic on its own, cells become sensitive to genomic hmdU when the uracil glycosylase SMUG1 hydrolyzes the base to create an abasic site (Fugger *et al*., 2021; Raja and Van Houten, 2021; Taglialatela *et al*., 2021)

To separate the roles of replication fork protection and ssDNA gap suppression in the response to hmdU, we rescued both pathways by depleting SMARCAL1 and PRIMPOL, respectively. We found that PRIMPOL-mediated gap formation is responsible for hmdU sensitivity and that this was specifically due to gap formation at abasic sites generated by SMUG1. These results affirm previous studies that implicated PRIMPOL in repriming DNA synthesis at stalled forks encountering abasic sites (Garcia-Gomez *et al*., 2013; Taglialatela *et al*., 2021), while directly linking gap formation by PRIMPOL to hmdU toxicity.

We also observed hypersensitization of the fork protection/gap suppression-deficient cells to combined olaparib and hmdU treatment, likely because olaparib triggers PARP1 trapping at abasic sites (Fugger *et al*., 2021). Again, this hypersensitization to the dual treatment is abrogated by PRIMPOL depletion, implicating gap formation in the sensitization. Given that PRIMPOL-mediated gaps arise with heterozygosity of BRCA2, modulating genomic hmdU levels may be an effective strategy for cancer chemotherapy for HDR-proficient tumors with gap suppression defects.

In contrast to PRIMPOL knockdown, depleting SMARCAL1 has no discernible effects, indicating that defective fork protection is not a significant contributor to therapy response. Because the replicative helicase remains bound during nascent strand degradation (Kavlashvili *et al*., 2023), fork protection-deficient cells can readily restart stalled replication forks (Schlacher *et al*., 2011). Thus, fork collapse may be relatively rare in these cells.

### HDR status ultimately determines response to PARP inhibition and hmdU

While synthetic lethality of PARP inhibition in BRCA-deficient cells was originally attributed to HDR deficiency (Bryant *et al*., 2005; Farmer *et al*., 2005), recent reports have called this into question (Cong *et al*., 2021; Ding *et al*., 2016; Ray Chaudhuri *et al*., 2016). However, using isogenic cell lines, we find that the main determinant of olaparib sensitivity is the status of HDR. While fork protection/gap suppression mutants have a 22% lower LD50 for olaparib than wild-type cells, HDR deficient cells have a 10-fold reduced LD50. Neither PRIMPOL nor SMARCAL1 knockdown affects olaparib sensitivity of the HDR mutant cells. Similarly, neither rescued hmdU sensitivity in HDR-deficient cells. However, SMUG1 knockdown does rescue hmdU sensitivity of HDR mutant cells, implying that abasic sites not at PRIMPOL-generated gaps ultimately require HDR for repair. We surmise that the aberrant DNA structures generated from degraded nascent strands, unfilled ssDNA gaps, or abasic sites may devolve into DSBs, including in the next cell cycle (Simoneau *et al*., 2021), which then require HDR for their repair. Our results affirm the importance of screening for homologous recombination deficiency when determining therapeutic response to PARP inhibitors, and highlight the merit of using PARPi/hmdU combined therapy and ssDNA gap screening to target tumors with gap suppression defects.

### Limitations of this study

We have yet to identify a BRCA2 allele that only causes HDR deficiency. The HDR-deficient cells (*Brca2^Δ27/Δ27^*) are additionally deficient in both fork protection and gap suppression. To address this limitation, we have knocked down proteins involved in these two processes (SMARCAL1 and PRIMPOL, respectively), but found no rescue of olaparib or hmdU sensitivity in this mutant. While it would be preferable to have a *Brca2* separation-of-function mutation, it may be difficult to identify such a mutation: Many of the same BRCA2 domains are required for all three processes, due to the common biochemical activity of promoting RAD51 filament formation.

Another limitation is that the *Brca2^Δ27/Δ27^* cells are hypomorphic. However, *Brca2^null/null^* die early in embryogenesis, prior to the ability to establish cell cultures. Moreover, the HDR defect of *Brca2^Δ27/Δ27^* cells is quite substantial (∼10-fold).

## Acknowledgements

We thank Travis White, Matteo Ferrari, Darpan Medhi and other members of the Jasin lab for helpful suggestions and critical discussions. We are grateful for the expert technical support from the past and present members of the MSK core facilities, including Peter Romanienko, Willie Mark, Ning Fan, and Eric Rosiek. This work was supported by P30CA008748, P50CA247749, BCRF-21-079, and R35CA253174 (M.J.).

## Author contributions

P.X.L. and M.J. conceived the study. P.X.L. performed the experimental work with assistance from M.Z. and under the supervision of M.J. P.X.L. and M.J. wrote the manuscript.

## Declaration of interest

The authors declare no competing interests.

**STAR METHODS**

**KEY RESOURCES TABLE**

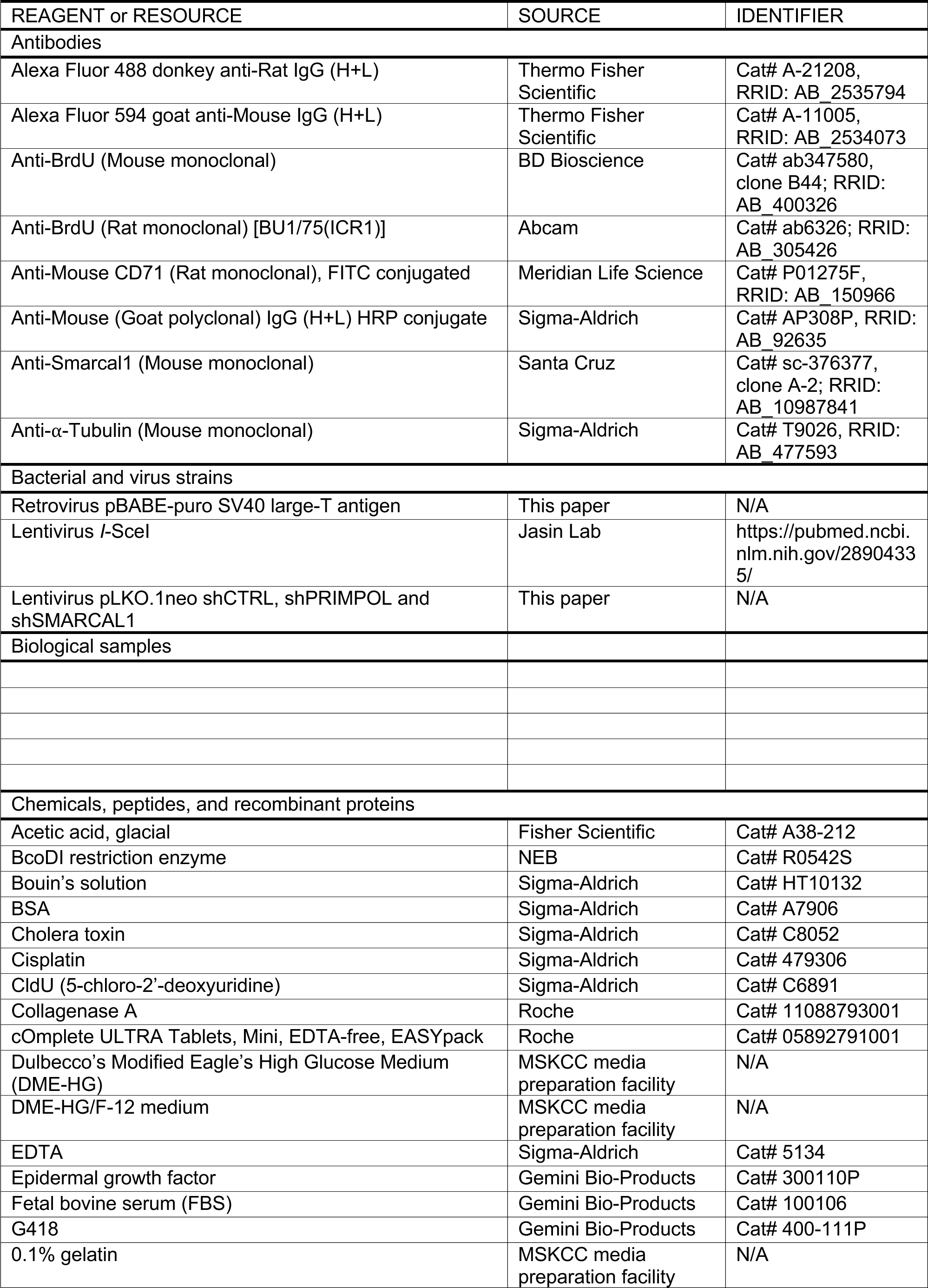
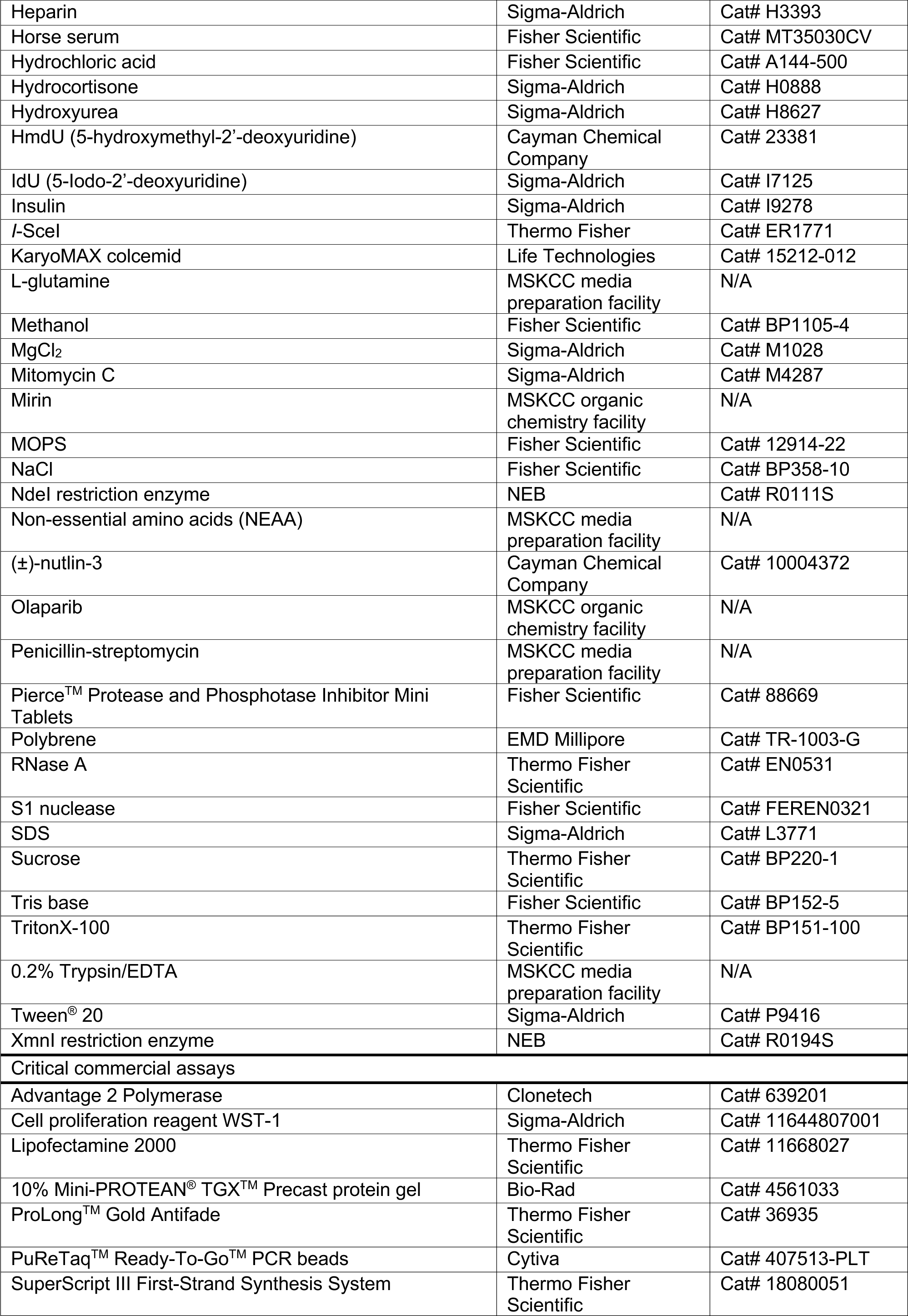
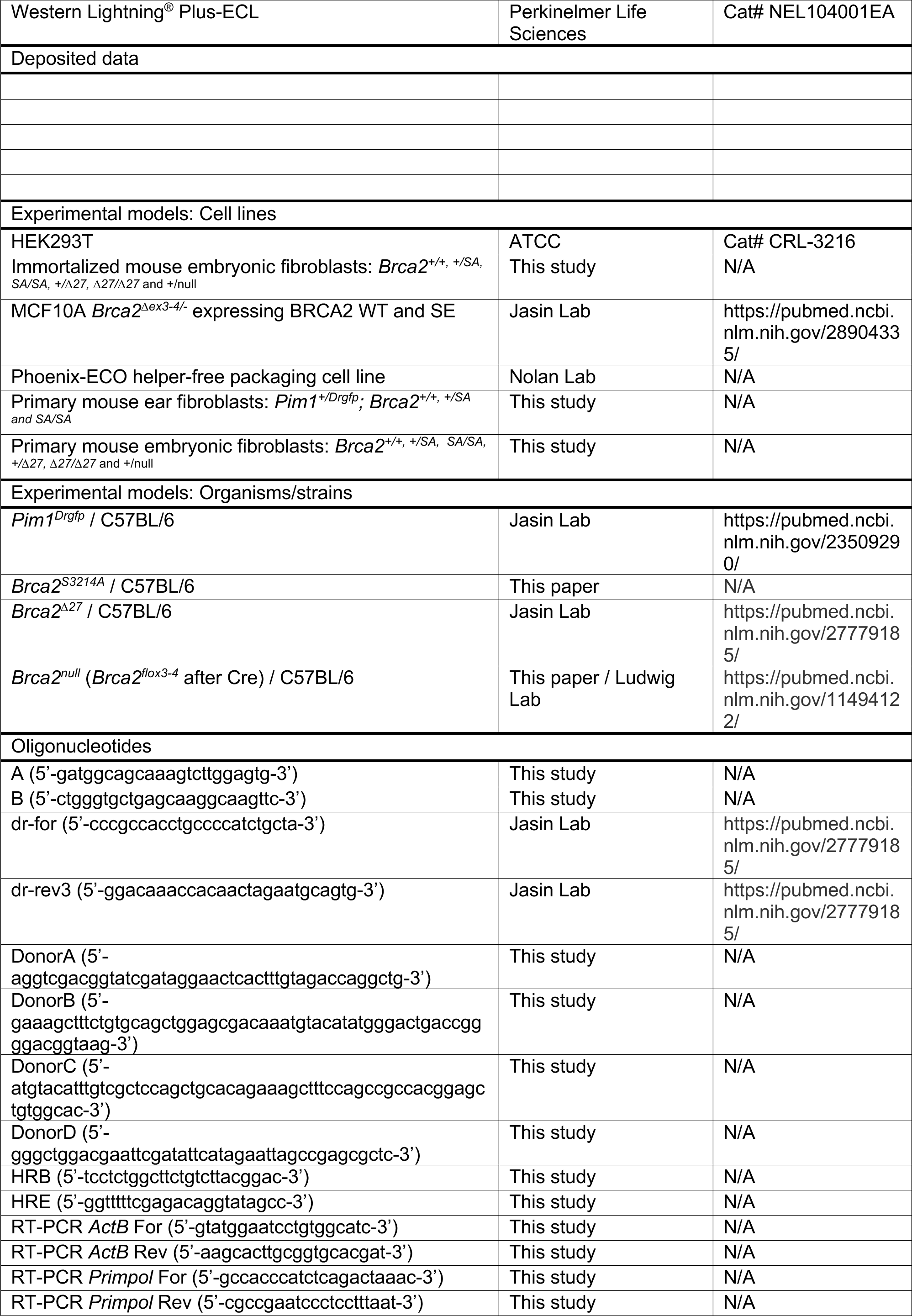
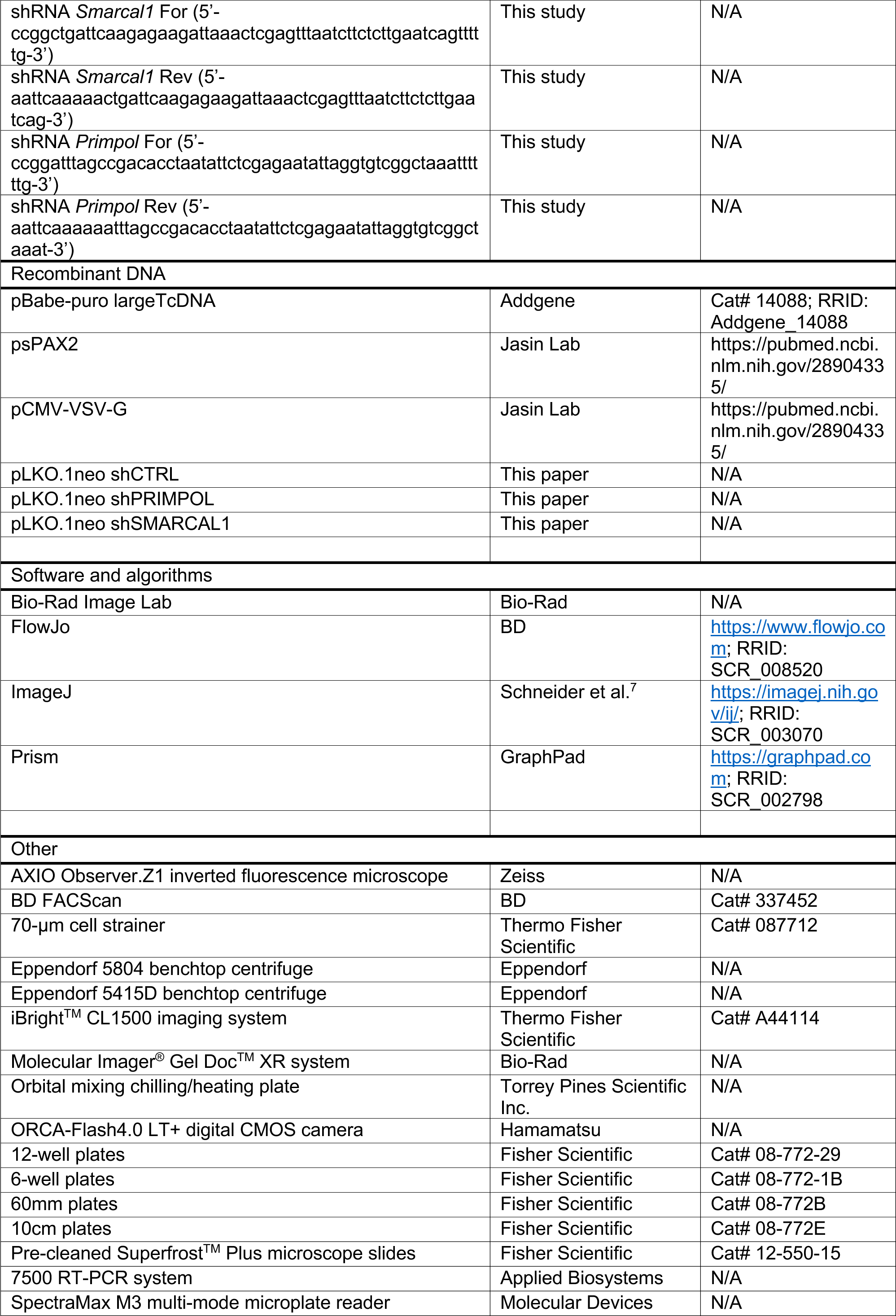
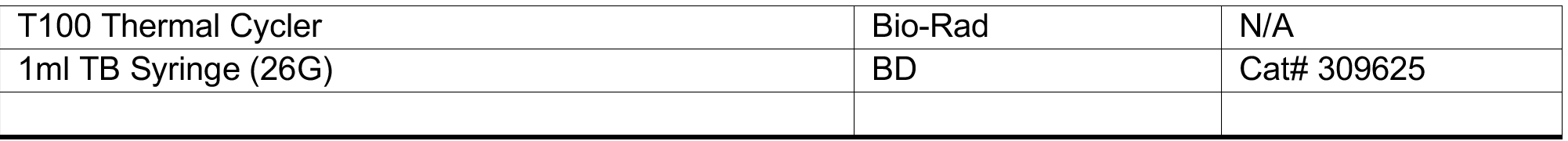

## RESOURCE AVAILABILITY

### Lead Contact

Further information and requests for resources and reagents should be directed to and will be fulfilled by the lead contact, Maria Jasin

### Materials availability

All materials generated in this study are available through lead contact upon request.

### Data and code availability

Raw data for all graphs and tables are provided in Supplemental Data File. Unprocessed blots, gels, microscopy, flow cytometry and immunohistochemistry images are available through lead contact upon request.

## EXPERIMENTAL MODEL AND SUBJECT DETAILS

### Mice

The *Brca2* S3214A mutation was generated in mice using TALENs (Cermak *et al*., 2011). The sequences for left and right TALEN recognition are 5’-CAGTCCCATCTGTACC and 5’-GAAAAGCCTTCTGTGCA, respectively (**Figure S1A**). The TCT codon for S3214 is centered within the spacer. To construct the donor template, the *Brca2* exon 27 region was amplified from mouse genomic DNA by PCR using Q5 High Fidelity DNA Polymerase (NEB) with primers containing the mutations. The 5’ homology was amplified using primers DonorA and DonorB and the 3’ homology was amplified using primers DonorC and DonorD. The S3214A mutation, within the underlined codon, and silent mutations are in bold.

DonorA: 5’-AGGTCGACGGTATCGATAGGAACTCACTTTGTAGACCAGGCTG.

DonorB: 5’-**G**AAAGC**T**TTCTGTGCAGCTGG<u>AG**C**</u>GACAAA**T**GTACA**T**ATGGGACTGACCGGGGACGGTAAG.

DonorC: 5’-**A**TGTAC**A**TTTGTC**<u>G</u>**<u>CT</u>CCAGCTGCACAGAA**A**GCTTT**C**CAGCCGCCACGGAGCTGTGGCAC.

DonorD: 5’-GGGCTGGACGAATTCGATATTCATAGAATTAGCCGAGCGCTC.

The PCR amplicons were then assembled using the Gibson Assembly Cloning Kit (NEB) to produce a plasmid containing the mutated *Brca2* exon 27 and flanking intronic sequences for total homology of 2101 bp containing 5 single bp changes (**Figure S1A**).

Approximately 2 pl total of mRNA for each TALEN (45 ng/µl) and donor plasmid containing the mutant exon 27 homology (15 ng/µl) was injected into the pronucleus of mouse zygotes. Injected zygotes were transferred to pseudo-pregnant recipient mice and 46 pups were produced. To identify founder mice, IN-IN PCR was performed on tail tip DNA using primers A (5’-GATGGCAGCAAAGTCTTGGAGTG) and B (5’-CTGGGTGCTGAGCAAGGCAAGTTC). Digestion with NdeI showed that 3 out of 46 mice had this site. These mice were further characterized with, 5’ OUT-IN PCR and 3’ IN-OUT PCR. Primers for 5’ OUT-IN PCR were HRE (5’-GGTTTTTCGAGACAGGTATAGCC) and primer B; likewise, primers for 3’ IN-OUT PCR were primers A and HRB (5’-TCCTCTGGCTTCTGTCTTACGGAC). Digestion of these PCR products confirmed the presence of the NdeI site and also the XmnI site. Sequencing confirmed that all 5 intended mutations, including S3214A were properly incorporated in two founders. Progeny from these founders were further backcrossed for 8 generations to obtain the S3214A mutation on a C57BL/6 background. Subsequent genotyping was done using the 5’ OUT-IN PCR primers and NdeI digestion.

### Cell culture

Mouse embryonic fibroblasts were cultured in Dulbecco’s Modified Eagle’s High Glucose Medium (DME-HG), supplemented with 10% FBS, 1% penicillin-streptomycin, 1% non-essential amino acids (NEAA) and 1% L-glutamine. MCF10A WT and S3291E cells (Feng and Jasin, 2017) were cultured in DME-HG/F-12 medium supplemented with 5% horse serum, 20 ng/ml epidermal growth factor, 0.5 mg/ml hydrocortisone, 100 ng/ml cholera toxin, 10 µg/ml insulin and 1% penicillin-streptomycin.

### Cell line generation and immortalization

For primary ear fibroblast isolation, ear tips were cut from 6- to 8-week-old mice, minced with a razor blade, and dissociated in 5 ml serum-free DMEM medium containing 2 mg/ml collagenase A (Roche) and 1X Pen-Strep on a 37°C shaker for 3 hr. The dissociated tissues were passed through a 70-μm strainer, pelleted by centrifugation at 400 g for 8 min, and plated to 60 mm dishes for 3-5 days before passing for experiments.

For primary mouse embryonic fibroblast isolation, E13.5 mouse embryos were separated from the placenta and embryonic sac, followed by removal of the head and liver, then minced, and dissociated in 0.05% trypsin at 37°C for 45 min in a 60-mm dish. The dissociated cells were resuspended by pipetting, pelleted at 400 g for 10 min, and plated to a 60-mm dish for 3-5 days before passing for experiments or immortalization.

To immortalize the primary MEFs, 5 X 10^5^ cells were seeded in a 6-well plate coated with 0.1% gelatin. The next day, cells were incubated with media containing polybrene and infected with retrovirus carrying the pBABE-puro SV40 large-T antigen construct (Zhao *et al*., 2003). Cells were centrifuged at 1,500 rpm for 30 min and incubated with the virus overnight. Then, cells were trypsinized and split into two 10-cm dishes coated with 0.1% gelatin. The following day, 10µM (±)-Nutlin-3 was added to one dish to select for p53-suppressed cells. After 3-4 days, media was removed, fresh (±)-Nutlin-3 were added and cells were allowed to grow for another 3-4 days. Then, the immortalized MEFs were passaged for freezing and for experiments.

## METHOD DETAILS

### DR-GFP and site-loss PCR assays

To measure HDR, 2 x 10^5^ primary ear fibroblast cells were plated in a 6-well dish and the next day infected with I-SceI lentivirus (Feng and Jasin, 2017). The virus was removed 24 hr later and fresh media was added to the cells. The percent GFP-positive cells were analyzed 48 hr later by flow cytometry. For the I-SceI site-loss assay (Kass *et al*., 2016), the remaining cells were pelleted and subjected to genomic DNA extraction. The sequence spanning the I-SceI cleavage site in the SceGFP gene was amplified from 150 ng of genomic DNA using Advantage 2 Polymerase (Clontech) with primers dr-for (5’-CCCGCCACCTGCCCCATCTGCTA) and dr- rev3 (5’-GGACAAACCACAACTAGAATGCAGTG). The PCR products were aliquoted in half and digested with 8 U I-SceI (Thermo Fisher) or buffer-only control *in vitro* at 37°C overnight and separated on a 1% agarose gel. Band intensities were quantified using Bio-Rad Image Lab software. The intensity of the uncut band (1.1 kb) was divided by the sum of the uncut (1.1 kb) plus the large cut band (0.9 kb; accounted for the size difference with a correction factor of 1.2) to calculate % I-SceI site loss. Cells with less than 20% site loss, which generally also showed lower % GFP positive signal, were considered having inefficient DNA cutting due to low lentiviral infection and were excluded from the analysis.

### Mitomycin C sensitivity

As previously described (Kass *et al*., 2013), mitomycin C (MMC, Sigma) was dissolved in sterile saline solution on the day of injection to yield 0.5 mg/ml solution. Mice between 6 to 10 weeks old were weighed and the amount of MMC solution to inject to achieve 3.5 mg/kg was calculated. MMC solution was delivered via a 26- gauge needle into the intraperitoneal region. Mice were monitored daily and euthanized when they had reached endpoint criteria (wasting, hunching, lethargy).

### DNA fiber assays

Primary MEFs between passage 2 to 5 were seeded in a 12-well plate at a density of 10^5^ cells per well. Immortalized MEFs and MCF10A cells were seeded in the same format as well. The experiments were performed as previously described (Quinet *et al*., 2017). The next day, they were pulse-labeled with 50 µM 5- iodo-2’-deoxyuridine (IdU; Sigma) for 20 min at 37°C. After IdU treatment, the cells were washed four to five times with warm media (DME-HG for MEFs and DME-HG/F12 for MCF10A) and pulse-labeled with 100 µM 5- chloro-2’-deoxyuridine (CldU; Sigma) for 20 min at 37°C. The cells were again washed with warm media and then either trypsinized (untreated control) or treated with 4 mM hydroxyurea (HU; Sigma), 2 hr for MEFs and 5 hr for MCF10A, followed by trypsin harvest. To inhibit the MRE11 nuclease, cells were treated with 50 µM mirin (MSKCC Organic Chemistry Core Facility) during pulse-labeling with IdU and CldU, as well as during the subsequent HU treatment. The pelleted cells were then resuspended in 1xPBS at a concentration of 10^6^ cells/ml. Two microliters of the cell suspension were pipetted onto pre-cleaned superfrost plus microscope slides (Thermo Fisher) and lysed with 8 µl spreading buffer (0.5% SDS, 200 mM Tris-HCl pH 7.4, 50 mM EDTA) for 6 min. The slides were then tilted to about 30° angle relative to horizontal allowing the cell lysis buffer mixture to flow down and spread the DNA fibers. Following spreading, the slides were air-dried for at least 20 min before fixing with ice cold fixative solution (75% methanol, 25% glacial acetic acid) for 4 min and storing in 4°C overnight or up 3 days. The DNA fibers were then denatured in 2.5 M HCl solution for 1 hr at room temperature and blocked with 5% bovine serum albumin (BSA; Sigma) in 1xPBS for 45 min at 37°C. The slides were incubated with 45 µl of 1:100 mouse anti-BrdU (BD Bioscience) and 1:100 rat anti-BrdU (Abcam) antibodies in 1% BSA, 0.05% Tween-20 in 1xPBS for 1.5 hr at room temperature in a humidified chamber. The slides were then stained with 45 µl of 1:100 anti-mouse Alexa Fluor 594 (Thermo Fisher) and 1:100 anti-rat Alexa Flour 488 (Thermo Fisher) in 1% BSA, 0.05% Tween-20 in 1xPBS for 1 hr at room temperature in a dark humidified chamber. Following secondary antibody staining, the slides were mounted in Prolong Gold Antifade (Thermo Fisher) and kept at 4°C until ready for imaging. Imaging of fibers was carried out on an AXIO Observer.Z1 inverted fluorescence microscope (Zeiss) with ORCA-Flash4.0 LT+ digital CMOS camera (Hamamatsu) at 63x magnification. Analysis was performed using imageJ software. At least 200 individual fibers were measured per experimental condition. The ratio of the lengths of adjacent IdU and CldU tracts were calculated and presented in a dot plot.

### S1 nuclease DNA fiber assay

Primary MEFs between passage 2 to 5 were seeded in a 12-well plate at a density of 2x10^5^ cells per well. Immortalized MEFs and MCF10A cells were seeded in the same format as well. The experiments were performed as previously described (Quinet *et al*., 2017) but with slight modifications. The next day, they were pulse-labeled with 50 µM IdU (Sigma) for 20 min at 37°C. After IdU treatment, the cells were washed four to five times with warm DMEM and chased with 100 µM CldU (Sigma) and HU (0.1 mM for MEFs and 0.5 mM for MCF10A) for 2 hr or 100 µM CldU and 30 µM olaparib for 1 hr at 37°C. Then, cells were trypsinized and split into two Eppendorf tubes coated with 2% BSA in PBS. Cells were centrifuged at 200 *g* for 5 min. After washing with 1xPBS, cells were permeabilized with 1ml CSK100 buffer (100 mM NaCl, 10 mM MOPS pH7, 3 mM MgCl_2_, 300 mM sucrose, 0.5% Triton X-100) at room temperature for 5 min. Cells were then centrifuged at 5000 *g* for 5 min to obtain a nuclear pellet. The pellet was washed with 1xPBS and treated with S1 nuclease (Thermo Fisher) at 37°C for 30 min with occasional tapping at 10-minute intervals. Following S1 nuclease treatment, the nuclear pellet was resuspended in 1 x PBS, spread on microscope slides and stained using the same protocol as the DNA fiber assay. Imaging of fibers and analysis were also done using the same protocol as the DNA fiber assay.

### shRNA knockdown

Constructs for shRNA targeting *Primpol* (5’- ATTTAGCCGACACCTAATATT) *Smarcal1* (5’- CTGATTCAAGAGAAGATTAAA) and *Smug1* (5’- AACTATGTGACTCGCTACT) were designed and inserted into pLKO.1neo plasmid (Addgene #13425). Lentiviruses were generated using a standard protocol (Feng and Jasin, 2017). Immortalized MEFs were infected with the lentivirus and cells stably expressing shRNA were selected using 200 µg/ml G418.

### RT-PCR

Total RNA was isolated using the conventional Trizol method and cDNA library was generated using the SuperScript^TM^ III First Strand Synthesis kit. Knockdown efficiency was determined using qRT-PCR with primers 5’-GCCACCCATCTCAGACTAAAC and 5’-CGCCGAATCCCTCCTTTAAT for *Primpol* amplification and primers 5’-GTATGGAATCCTGTGGCATC and 5’-AAGCACTTGCGGTGCACGAT for β-actin amplification as an internal control. The experiment was run in Applied Biosystems 7500 RT-PCR system in triplicates and analyzed using the relative quantification (RQ or 2^-ΔΔCT^) method.

### Western blotting

Equal amounts of protein samples were heated at 95°C for 5 mins, run on a 10% Mini-PROTEAN^®^ TGX^TM^ protein gel (Bio-Rad) and then transferred to a PVDF membrane. The membrane was blocked in 5% non-fat dry milk in PBST for 1 hr and incubated in SMARCAL1 (1:500; sc-376377, Santa Cruz Biotechnology) or tubulin (1:10,000; T9026, Sigma) primary antibodies at 4°C overnight, followed by incubation in HRP-linked anti-mouse IgG (1:10,000; Sigma) secondary antibody at room temperature for 1 hr.

### Metaphase spreads

The experiments were performed as previously described (Weiss *et al*., 2000). Briefly, primary MEFs between passage 2 to 5 were seeded in a 60-mm dish at a density of 5 x 10^5^ cells. The next day, they were treated with 4 mM HU (or fresh media for untreated control) for 2 hr. After HU removal, the cells were treated with 0.1 µg/ml colcemid (Invitrogen) for 1-4 hr until sufficient rounded (metaphase arrested) cells had accumulated. The cells were then trypsinized and collected in a 15-ml conical tube. The cells were incubated in 6 ml pre-warmed hypotonic solution (75 mM KCl in H_2_O) for 6 min in a 37°C water bath. Following hypotonic swelling, 1 ml fresh ice cold fixative solution (75% methanol, 25% glacial acetic acid) was added and mixed gently. Cells were centrifuged at 250 *g* for 5 min and resuspended in 5 ml fresh cold fixative solution. Cells were incubated on ice for 1 hr and washed with fresh cold fixative solution 3 times before resuspended in fresh cold fixative solution and spotted on superfrost plus microscope slide (Thermo Fisher). Slides were allowed to air dry overnight before being mounted in Prolong Gold Antifade with DAPI (Thermo Fisher). Imaging of metaphase spreads was carried out on an AXIO Observer.Z1 inverted fluorescence microscope (Zeiss) with ORCA-Flash4.0 LT+ digital CMOS camera (Hamamatsu) at 100x magnification. Analysis was performed using imageJ software (cell counter plugin). About 45-50 spreads were quantified for each experimental condition. Three repeats were done for each genotype and the total quantification was compiled together.

### Testis histology

Testes from 2- to 4-month-old mice were collected and fixed overnight in Bouin’s fixative solution, embedded in paraffin and sectioned. Sections were stained with hematoxylin and eosin successively. Images were acquired with a Mirax scanner.

### Micronucleus assay

The micronucleus assay was performed as previously described (Dertinger *et al*., 1996). Briefly, three drops of peripheral blood were collected from 1.5− to 4-month-old mice via submandibular vein puncture. They were stored in Eppendorf tubes containing 200 µl heparin solution before fixing in 100% cold methanol in -80°C overnight. After washing and centrifugation, cell pellets were resuspended in wash buffer containing CD71− FITC antibody (Meridian Life Science) and RNaseA (Thermo Fisher). Cells were stained shielded from light in 4°C for 45 min. Then, cells were washed, pelleted and resuspended in wash buffer containing propidium iodide (PI; Sigma). At least 100,000 total erythrocytes from each of these samples were analyzed using BD FACScan with the accompanying CellQuest software. FL1 and FL3 channels were used to detect CD71 and PI respectively. The overlapping peaks were manually compensated in CellQuest. Using Flowjo, normochromic erythrocytes (NCE) were quantified by gating for CD71^−^PI^−^ population, whereas micronucleated NCE (MN-NCE) were quantified by gating for CD71^−^PI^+^ population. Percent MN-NCE was calculated from the number of MN-NCE cells divided by the total number of NCE and MN-NCE cells.

### Tumor monitoring

Mice were monitored weekly for tumor development (Billing *et al*., 2018). Upon detection of humane endpoints such as palpable mass, distended abdomen, kyphosis or skin lesion, the mice were euthanized with CO_2_ and necropsy was performed. Organs with suspected hyperplasia or tumor formation were harvested and fixed at 4°C overnight in 10% buffered formalin, followed by dehydration with 70% ethanol the next day. Tissues were then embedded, sectioned and stained with hematoxylin and eosin for histopathological evaluation. Overall survival and tumor-free Kaplan-Meier survival curves were plotted in GraphPad Prism.

### Genotoxin sensitivity assays

For each experiment, two independently derived immortalized MEFs from each *Brca2* wildtype and mutant genotype between passage 10 and 30 were seeded in triplicates in a gelatinized 96-well plate at a density of 2,500 cells/well. The next day, cells were treated with HU, cisplatin (Sigma), olaparib (MSKCC Organic Chemistry Core Facility) or olaparib with 5-hydroxymethyl-2’-deoxyuridine (hmdU; Cedarlane Labs) at various concentrations. For sensitivity assay with HU, cells were treated with 0, 20, 50, 100, 200, or 500 µM HU for 24 hr before changing to fresh media and counted 5 days later. For sensitivity assay with cisplatin, cells were treated with 0, 10, 25, 50, 200 or 500 nM cisplatin for 6 days with fresh cisplatin given on the third day. For sensitivity assay with olaparib, cells were treated with 0, 0.5, 1, 2.5, 5, or 7.5 µM olaparib with or without 0.5 µM hmdU for 6 days with fresh olaparib given on the third day. Conversely, cells were treated with 0, 0.2, 0.5, 1, 2, or 5 µM hmdU with or with the addition of 0.1 µM olaparib for 6 days with fresh hmdU and olaparib given on the third day. At least two repeats were carried out for each experiment. To count cells after treatment, the cells were given 10 µl of cell proliferation reagent WST-1 (Sigma) per well and incubated for 1 hr at 37°C. Thereafter, the plate was placed in a SpectraMax M3 multi-mode microplate reader (Molecular Devices), shaken, and read for absorbance values at 450 nm (experimental) and 680 nm (reference) wavelengths. Data analysis was done in Prism 9 (GraphPad). All absorbance values were normalized to the values of untreated cells and plotted in XY graphs against treatment concentrations in log scale. LD50s were estimated from non-linear regression curve fit analyses.

## QUANTIFICATION AND STATISTICAL ANALYSIS

Statistical analyses were performed using GraphPad Prism (Version 9.0). Statistical parameters and tests are reported in the figures and corresponding figure legends. To reiterate, all comparisons in DNA fiber assays were determined using Mann-Whitney U test; mouse Kaplan-Meier survival curve comparisons were conducted using Log-rank test; comparisons for DR-GFP, testis size, micronucleus levels, chromosome aberrations in metaphase spreads, LD50 values and single dose survival assays were performed using unpaired Student’s t test. Comparisons for dose-response curves were done using two-way ANOVA test. In all cases, ns, not significant (p>0.05); *, p<0.05; **, p<0.01; ***, p<0.001; and ****, p<0.0001.

---

**Figure S1.**
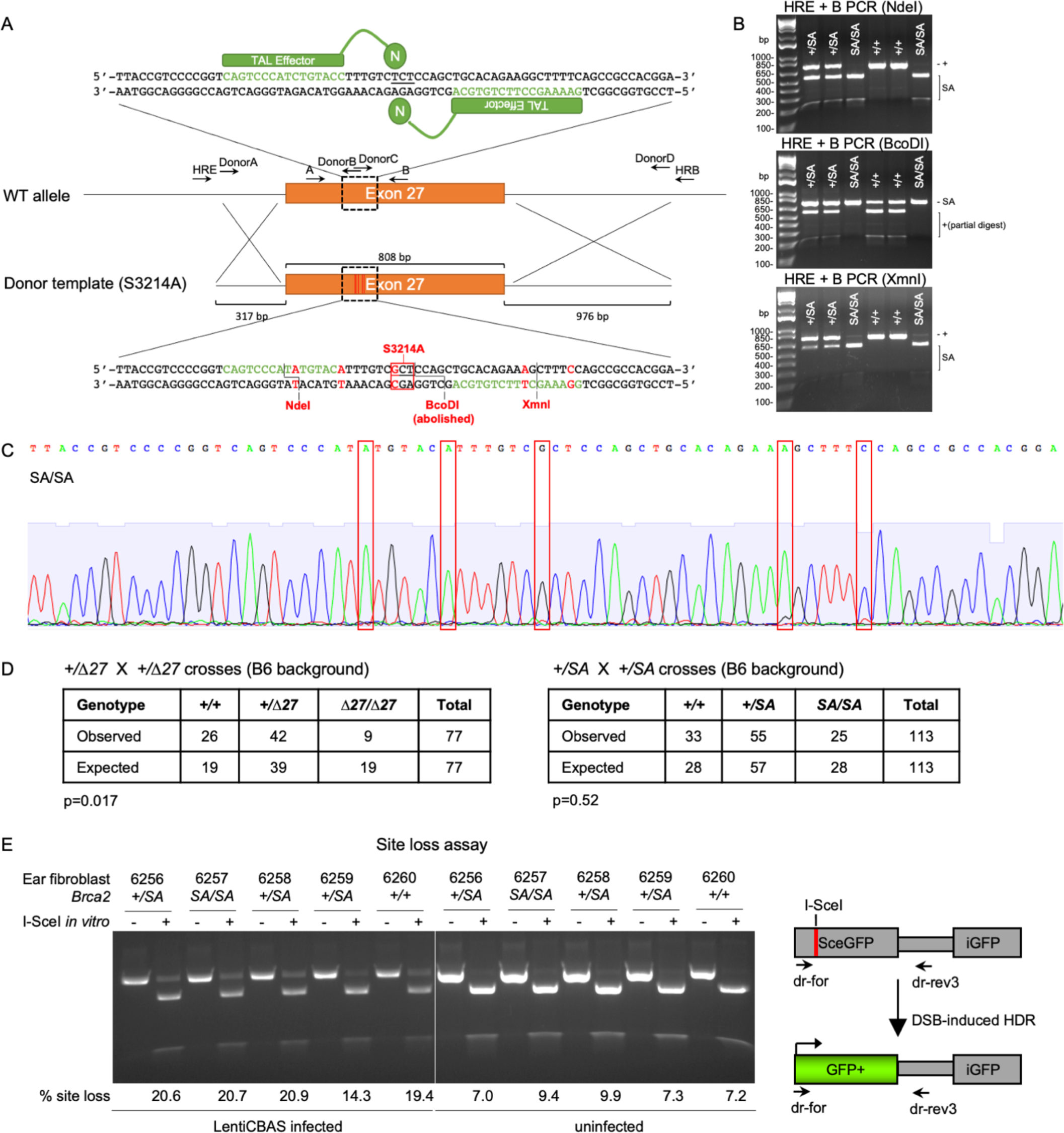
Generation of the *Brca2^S3214A^* allele in mice and I-SceI site loss for the DR-GFP assay. **A.** Strategy to knock-in the S3214A mutation into exon 27 of the endogenous *Brca2* allele. TAL effectors were designed to recognize 16-nucleotide sequences (green) flanking the S3214 TCT codon to make a double-strand break (DSB) with the fused nuclease (N). The DSB was repaired from a dsDNA donor template which contains a 5’ homology arm, the entire *Brca2* exon 27 sequence containing five single nucleotide substitutions (red), and a 3’ homology arm. The five substitutions include the intended serine to alanine mutation (red box), which abolished a BcoDI restriction site, and two silent mutations in each of the TAL effector recognition sequences to prevent donor template cutting and to introduce two new restriction sites (NdeI and XmnI) for genotyping. Primer locations were approximated on WT allele. **B.** Genotyping the S3214A allele in mice. The PCR product generated from the *Brca2^S3214A^* allele in mice is fully digested by the NdeI and XmnI restriction enzymes, while resistant to BcoDI. The PCR product from the wild-type allele is resistant to NdeI and XmnI, although only partially digested by BcoDI. **C.** Sanger sequencing chromatogram of a *Brca2^SA/SA^* mouse verifying the five single nucleotide substitutions introduced by gene editing (red boxes). **D.** Heterozygous crosses of *Brca2^+/Δ27^*and *Brca2^+/SA^* mice. *Brca2^SA/SA^* mice are observed at the normal Mendelian ratio, whereas *Brca2^Δ27/Δ27^* mice are underrepresented. Chi-square tests were performed and the p-values are shown below. **E.** I-SceI site loss assay to quantify percent site loss as a measure of repaired DSBs (a combination of HDR and NHEJ). Primary ear fibroblasts from one of the 5 sets of DR-GFP experiments were lysed to extract genomic DNA. The GFP sequence flanking the I-SceI site was amplified and digested *in vitro* with I-SceI. The PCR product is cleaved *in vitro* if a DSB was not induced and/or visibly repaired, but not cleaved *in vitro* if repair by HDR or NHEJ *in vivo* leads to I-SceI site loss. Percent site loss is calculated by dividing the intensity of uncleaved band with the total intensity of cleaved and uncleaved bands. Low % site loss may indicate inefficient lentivirus infection, and the data for DR-GFP for that cell line (14.3%) was excluded. The uninfected cells showed an average of 8.2% site loss, indicating incomplete *in vitro* I-SceI digestion, which is treated as background.

**Figure S2.**
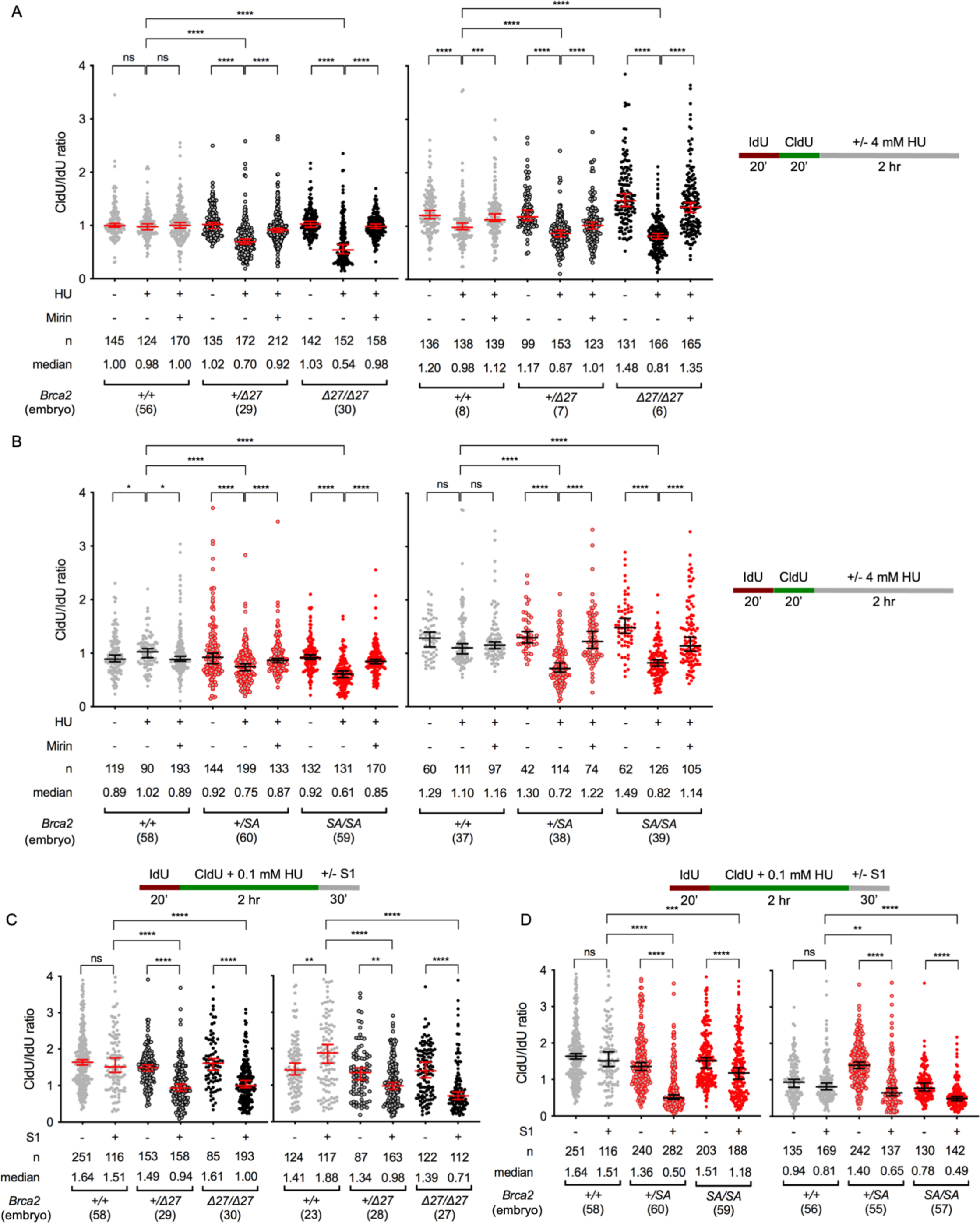
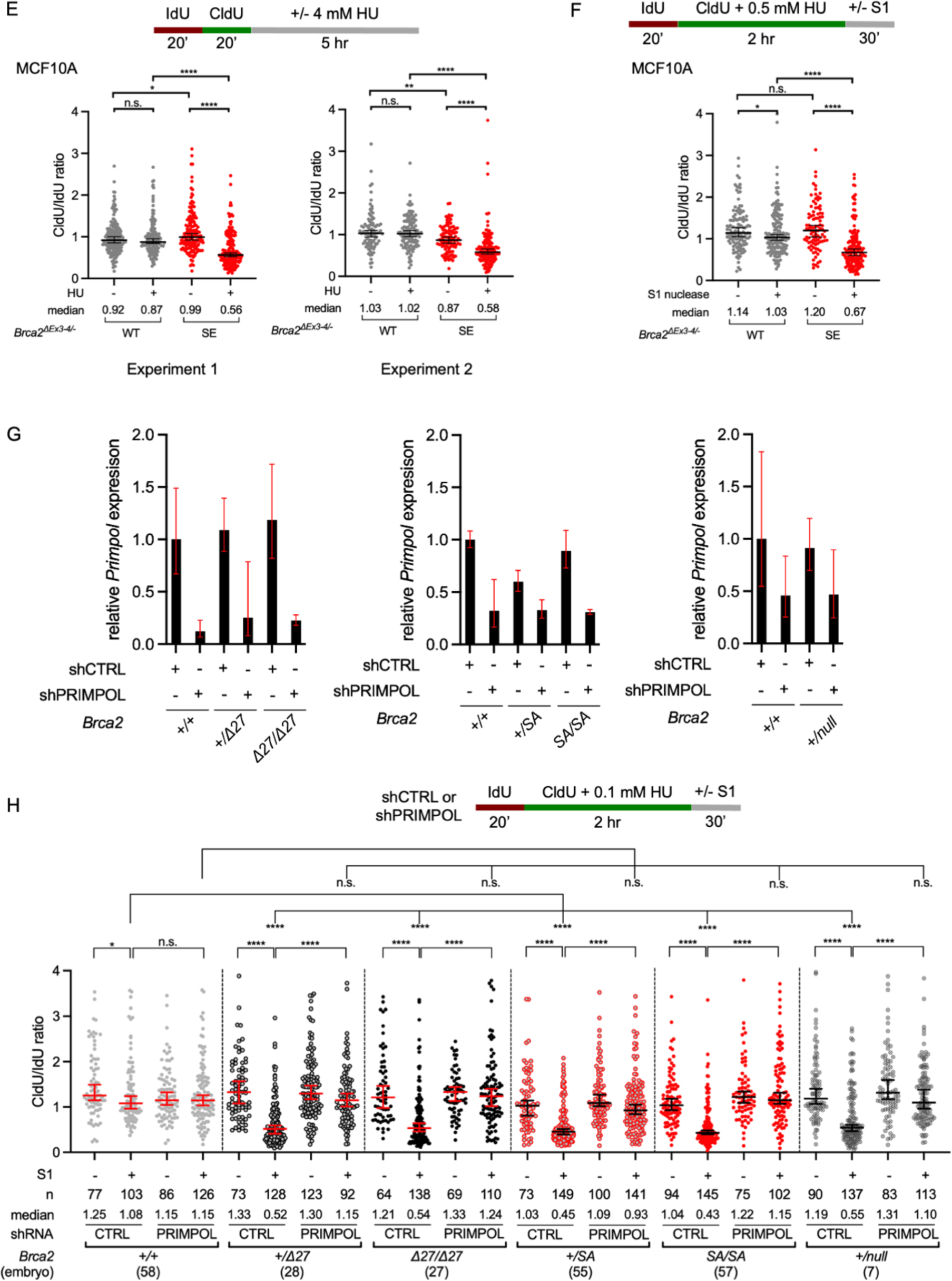
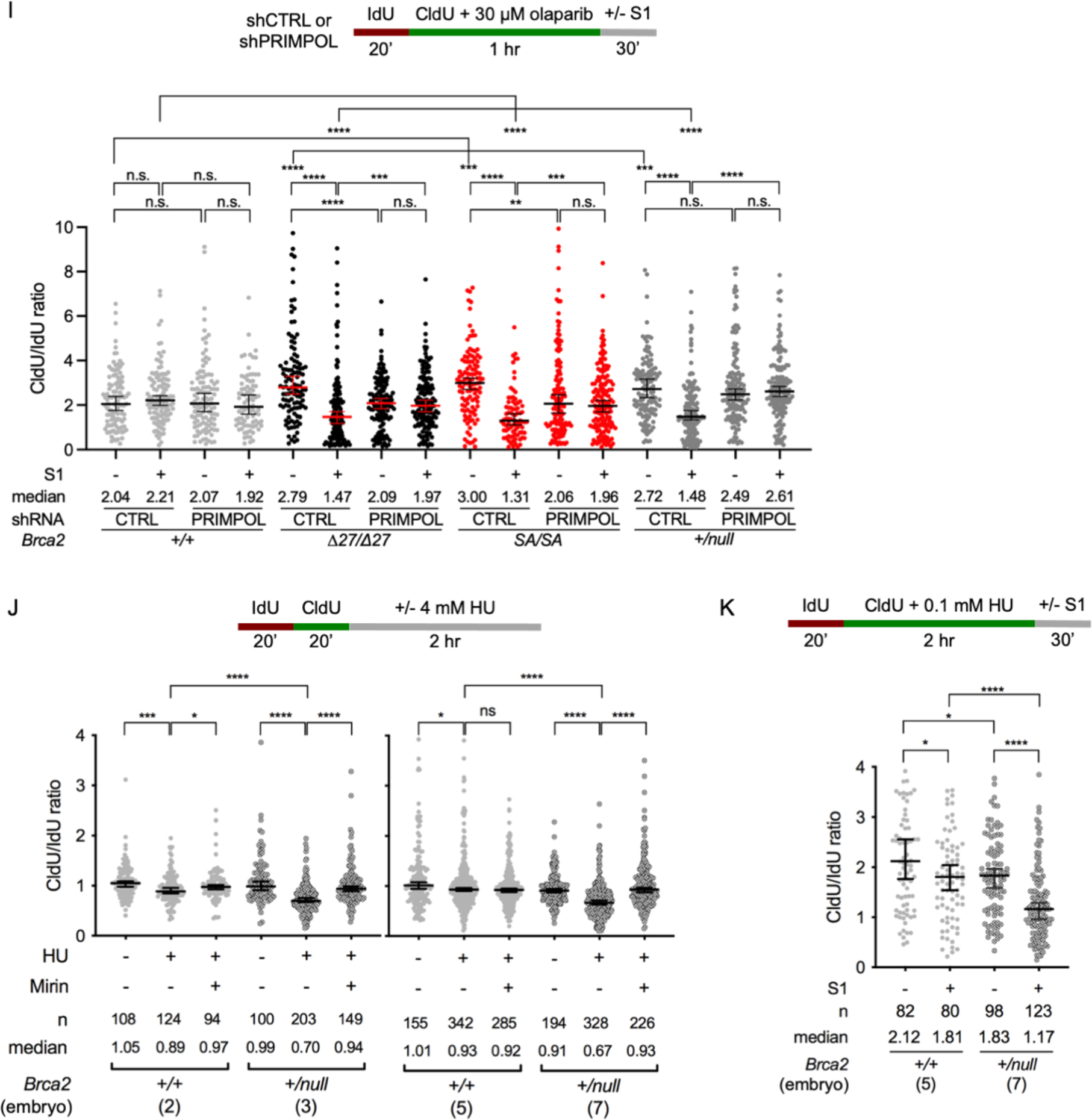
BRCA2 C-terminal RAD51-interacting site is required for stalled fork protection and gap suppression. **A-D, H-K.** Repeat experiments for the DNA fiber assays shown in the main text using different embryos to derive MEFs. Numbers in parentheses refer to the embryos from which primary MEF were derived. n, number of IdU+CldU tracks analyzed. Statistical analysis, Mann-Whitney U test (median ± 95% confidence intervals). ns, not significant; *, p<0.05; **, p<0.01; ***, p<0.001; ****, p<0.0001. **A,B.** Degradation of stalled forks with deletion (Δ27) (**A**) or mutation (SA) (**B**) of the BRCA2 C-terminal RAD51- interacting site in mouse cells. Repeat experiments are shown for the DNA fiber assays in Figure 1D using primary MEFs. **C,D.** Gap suppression defects with deletion (Δ27) (**C**) or mutation (SA) (**D**) of the BRCA2 C-terminal RAD51- interacting site in mouse cells. Repeat experiments are shown for the DNA fiber assays in Figure 2A using primary MEFs. **E.** Degradation of stalled forks with BRCA2 S3291E (SE) mutation in immortalized MCF10A human mammary epithelial cells. **F.** Gap suppression defects with with BRCA2 S3291E (SE) mutation in immortalized MCF10A human mammary epithelial cells. A repeat experiment is shown for the DNA fiber assays in Figure 2B. **G.** Knockdown of PRIMPOL in the immortalized MEFs. Expression of *Primpol* mRNA relative to β-actin mRNA was determined by qPCR. Error bars denote relative quantification (RQ) min and max values. **H.** PRIMPOL knockdown suppresses gap formation caused by HU in *Brca2* mutant cells. Repeat experiments are shown for the DNA fiber assays in Figure 2D,**2G** using immortalized MEFs. **I.** PRIMPOL knockdown suppresses gap formation caused by olaparib in *Brca2* mutant cells. **J.** BRCA2 haploinsufficiency for the protection of stalled fork in mouse cells. Repeat experiments are shown for the DNA fiber assays in Figure 2E using in primary MEFs. **K.** BRCA2 haploinsufficiency for the gap suppression in mouse cells. A repeat experiment is shown for the DNA fiber assays in Figure 2F using in primary MEFs.

**Figure S3.**
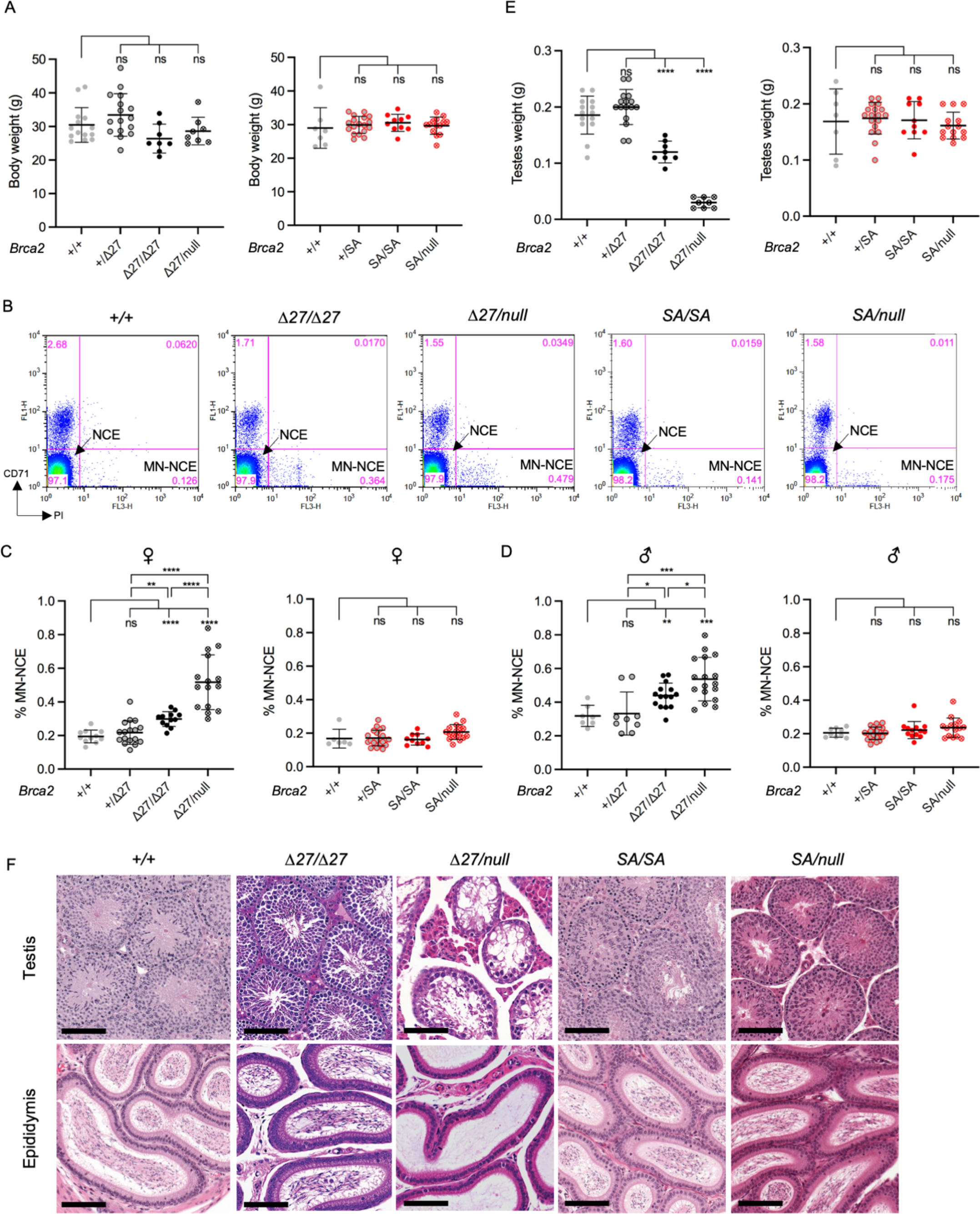
BRCA2-mediated replication fork protection and gap suppression are not required for genomic integrity or normal testis development in mice. **A.** Body weight plots are not significantly altered in any of the *Brca2* mutants. Statistical analysis, unpaired Student’s t test (mean ± 1 SD). ns, not significant. **B.** Representative flow cytometry scatter plots for micronuclei (MN) detection in normochromic erythrocytes (NCEs). The fluorescence intensity of PI is on the x-axis and that of CD71 is on the y-axis. NCE cells are CD71^−^PI^−^ (lower left quadrant) and MN-NCE cells are CD71^−^PI^+^ (lower right quadrant). **C,D.** MN-NCEs are increased in HDR-deficient *Brca2^Δ27/Δ27^* and especially *Brca2^Δ27/null^* cells but not in cells only deficient for fork protection/gap suppression. Percent MN-NCE are shown separately for female (**C**) and male (**D**) mice. Statistical analysis, unpaired Student’s t test (mean ± 1 SD). ns, not significant; *, p<0.05; **, p< 0.01; ***, p<0.001; ****, p<0.0001. **E.** Testes weight are reduced in HDR-deficient *Brca2^Δ27/Δ27^* and especially *Brca2^Δ27/null^* mice. Statistical analysis, unpaired Student’s t test (mean ± 1 SD). ns, not significant; ****, p<0.0001. **F.** *Brca2^Δ27/null^*mice, but not other *Brca2* mutants, have perturbed spermatogenesis and absence of sperm in the epididymus. Representative hematoxylin and eosin sections of testis and caput epididymis are shown. With the exception of *Brca2^Δ27/null^*mice, all of the mutant mice had normal seminiferous tubules with the full range of spermatogenic stages, whereas *Brca2^Δ27/null^* mice had sparse spermatogonia and aberrant or absent meiotic stages. Scale bar, 100 µm.

**Figure S4.**
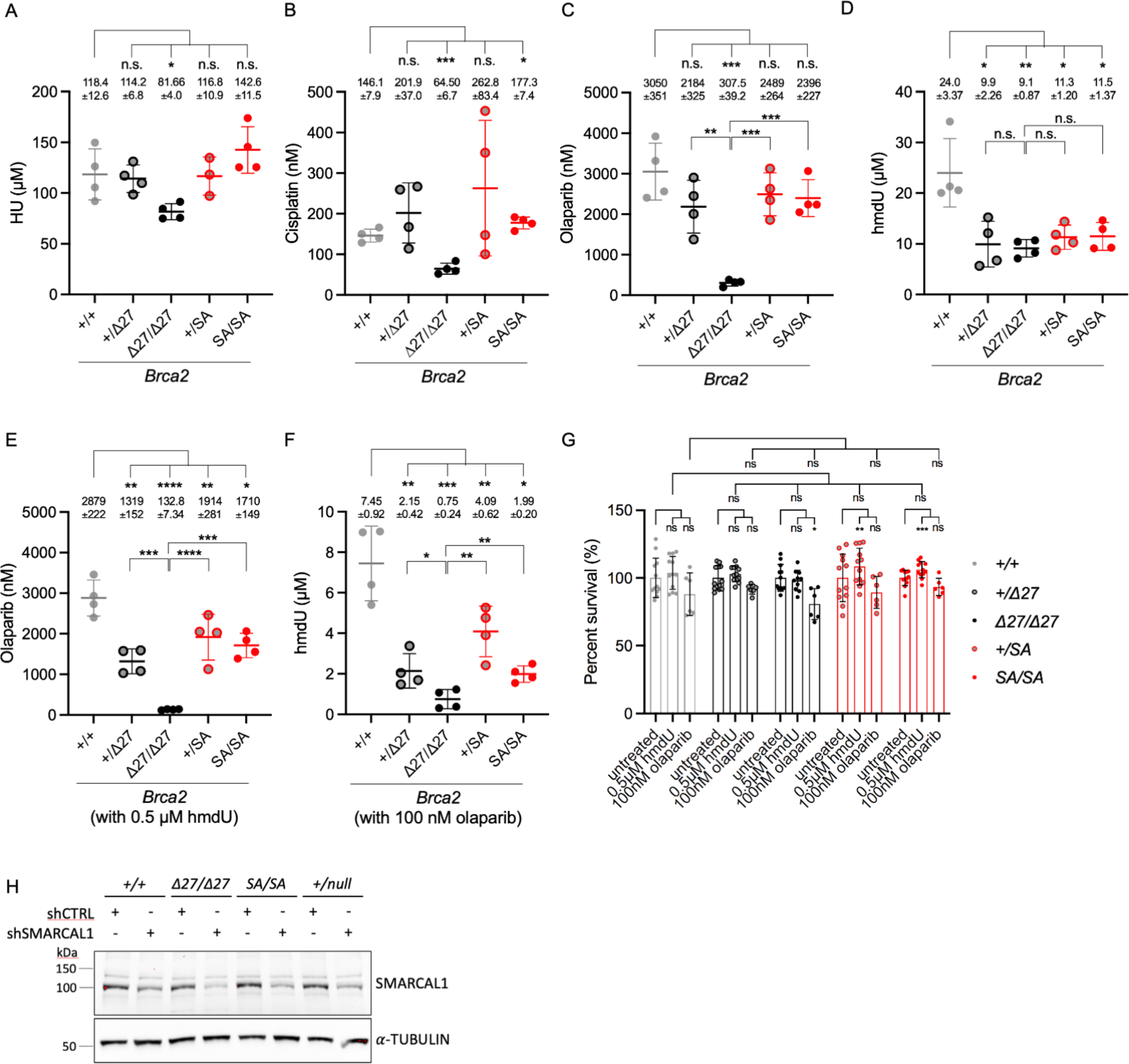
Genotoxin sensitivity is greatest in the absence of HDR, but is observed for hmdU in fork protection/gap suppression defective cells. **A-F.** The 50% inhibitory concentrations (LD50s) from non-linear regression curves of the sensitivity assays (Figure 4) using HU (**A**), cisplatin (**B**), olaparib (**C**), hmdU (**D**), olaparib with a fixed concentration of hmdU (**E**), or hmdU with a fixed concentration of olaparib (**F**). The average LD50 of the *Brca2^Δ27/Δ27^*immortalized MEFs was consistently lower than that of the wild-type cells. MEFs with only fork protection/gap suppression defects show milder or no sensitivity, except in the case of hmdU alone, in which the sensitivities of all *Brca2* mutants are more similar (**D**) and are exacerbated by olaparib (**E,F**). Statistical analysis, unpaired Student’s t test (mean ± 1 SD). ns = not significant; *, p<0.05; **, p< 0.01; ***, p<0.001; ****, p<0.0001. **G.** Cell survival for *Brca2* mutants after fixed, low-dose treatment of hmdU (Figures 4E**, S4E**) or olaparib (Figures 4F**, S4F**). Cell survival for all the *Brca2* mutants was not significantly different from wild type for either treatment. Statistical analysis, unpaired Student’s t test (mean ± 1 SD). ns = not significant; *, p<0.05; **, p< 0.01; ***, p<0.001. **H.** Western blot for SMARCAL1 in immortalized MEFs upon stable shRNA expression.

